# Direct binding of chromosome axis and cohesin complexes underlies meiotic chromosome architecture in fungi and plants

**DOI:** 10.64898/2026.01.19.700424

**Authors:** Francisco Mendez Diaz, Carolyn R. Milano, Jialu Huo, Andreas Hochwagen, Kevin D. Corbett

## Abstract

In prophase of meiosis I, the proteinaceous chromosome axis provides a scaffold for the compaction of chromosomes into a linear loop array, controls the formation of interhomolog crossovers, and finally becomes integrated into the synaptonemal complex after crossovers have formed. Despite its fundamental importance, how the proteins of the meiotic chromosome axis – meiotic HORMADs, axis core proteins, and cohesin complexes – self-assemble with one another is incompletely understood. In particular, it remains unknown how cohesin complexes interact with other axis components. Here, we combine genetics in *S. cerevisiae*, AlphaFold-based protein interaction screens, and biochemical assays to reveal that a conserved motif in the *S. cerevisiae* axis core protein Red1 – the cohesin-interacting motif or CIM – binds the C-terminal winged helix domain of cohesin’s meiosis-specific kleisin subunit Rec8. Disrupting the Red1 CIM specifically reduces Rec8-dependent Red1 association with meiotic chromosomes, and results in a strong spore viability defect. Finally, we find that the identified axis core-kleisin interaction is conserved across fungi and plants, but not in mammals, suggesting that different eukaryotic phyla use distinct protein-protein interfaces to assemble the chromosome axis.

## Introduction

All sexually-reproducing organisms undergo meiosis, a two-stage cell division program that produces haploid gametes from a diploid precursor cell (Zickler & Kleckner, 2023). During meiotic prophase I, the topoisomerase-like enzyme Spo11 introduces programmed double-strand DNA breaks (DSBs) along each chromosome (Zheng *et al*, 2025; Oger & Claeys Bouuaert, 2025). These breaks are repaired by the homologous recombination pathway using the homologous chromosome (termed the homolog) as a template, eventually leading to the formation of inter-homolog crossovers or chiasmata. Crossovers mediate the exchange of alleles between homologs to promote genetic diversity, and physically link each pair of homologs to ensure proper chromosome segregation in the ploidy-reducing meiosis I division. Following meiosis I, a second round of cell division (meiosis II) segregates replicated sister chromosomes to complete meiosis.

Successful meiotic recombination depends on a proteinaceous assembly called the chromosome axis, which organizes chromosomes into a linear array of chromatin loops, promotes Spo11-mediated DSB formation, and orchestrates inter-homolog crossover formation (Ito & Shinohara, 2022; Ur & Corbett, 2021). In most eukaryotes, including budding yeast (*Saccharomyces cerevisiae*), plants, and mammals, the chromosome axis comprises three conserved components: (1) HORMA-domain proteins (HORMADs; Hop1 in *S. cerevisiae*, ASY1 in *A. thaliana*, and HORMAD1/2 in *M. musculus*), (2) filamentous “axis core” proteins (Red1 in *S. cerevisiae*, ASY3/4 in *A. thaliana*, and SYCP2/3 in *M. musculus*), and (3) cohesin complexes containing a meiosis-specific kleisin subunit (Rec8 in *S. cerevisiae*, REC8(SYN1) in *A. thaliana*, and REC8/RAD21L in *M. musculus*). In general, HORMADs are thought to control DSB formation and repair (Dereli *et al*, 2024; Shin *et al*, 2010; Stanzione *et al*, 2016; Milano *et al*, 2024), the cohesin complex binds DNA and extrudes and/or constrains chromatin loops to form the basis of the axis (Nagasaka *et al*, 2023; GuérinThomas, 2025), and the axis core proteins provide a physical scaffold for assembly of HORMADs on the axis (West *et al*, 2019).

The architecture and interactions of axis core proteins and meiotic HORMADs is generally conserved across yeast, plants, and mammals. Meiotic HORMADs possess an N-terminal HORMA domain which can adopt an open/unbuckled conformation, or a closed conformation in which it is tightly bound to a “closure motif” peptide (West *et al*, 2018; Wang *et al*, 2023). HORMADs possess a closure motif at their C-terminus that is thought to bind the HORMA domain to generate an inactive “self-closed” state (Yang *et al*, 2020). Conversion to an open/unbuckled state by the AAA+ ATPase Pch2/TRIP13 enables the HORMA domain to bind a higher-affinity closure motif in an axis core protein, mediating its localization to the chromosome axis (Ye *et al*, 2017; West *et al*, 2018; Yang *et al*, 2020; Raina & Vader, 2020). Mutation of the closure motif in *S. cerevisiae* Red1 abolishes Hop1 binding and causes severe defects in meiotic recombination and homolog synapsis (Woltering *et al*, 2000), demonstrating the importance of axis core-HORMAD binding. Many HORMADs also possess a central “chromatin binding region” (CBR) with variable domain structure, which can independently bind DNA and/or chromatin (Rodriguez *et al*, 2025; Milano *et al*, 2024). The biological role of the HORMAD CBR is still unknown, but it may mediate enrichment of axis proteins at particular chromosome regions and/or play a signaling role during recombination (Milano *et al*, 2024).

Axis core proteins possess a conserved overall architecture with a folded N-terminal domain comprising tightly-packed armadillo-repeat (ARM) and pleckstrin homology (PH; also termed Spt16M-like) subdomains (missing in plant ASY3), a central HORMAD-binding closure motif, an extended disordered region, and a C-terminal coiled-coil domain (West *et al*, 2019; Feng *et al*, 2017). In *S. cerevisiae* Red1, the coiled-coil domain assembles into homotetramers, and these tetramers further assemble into filaments that could form a scaffold that stabilizes the chromosome axis (West *et al*, 2019). In plants and mammals, one axis core protein (ASY3 in plants, SYCP2 in mammals) forms 2:2 heterotetramers with a binding partner that contains only a short N-terminal disordered region and a C-terminal coiled-coil (ASY4 in plants, SYCP3 in mammals); these complexes further assemble into filaments as in *S. cerevisiae* Red1 (West *et al*, 2019; Syrjänen *et al*, 2014; Chambon *et al*, 2018).

Cohesin complexes are large multi-subunit assemblies with a pair of SMC ATPase subunits (in *S. cerevisiae*, Smc1 and Smc3), a “kleisin” subunit (in *S. cerevisiae*, Scc1 (Mcd1) in mitosis and Rec8 in meiosis), and a HEAT repeat subunit (in *S. cerevisiae*, Scc3 (Irr1)). Cohesin complexes play two roles in meiotic chromosome organization. One population of cohesin is deposited during pre-meiotic DNA replication, and maintains the cohesion (physical association) of replicated sister chromosomes (Eijpe *et al*, 2003; Srinivasan *et al*, 2020; Lengronne *et al*, 2006; Watanabe & Nurse, 1999). Another more dynamic population binds DNA and is thought to extrude and/or constrain large DNA loops (∼20 kb in *S. cerevisiae*; up to ∼1 Mb in *M. musculus*) to compact and organize meiotic chromosomes (Schalbetter *et al*, 2019; Patel *et al*, 2019; Alavattam *et al*, 2019). While the cohesive subpopulation comprises the four subunits described above, dynamic loop-extruding cohesin contains an additional two subunits (in *S. cerevisiae*, Scc2 and Scc4) (Davidson *et al*, 2019; Kim *et al*, 2019). In some organisms, there is a division of labor between cohesin subpopulations with different kleisin subunits: for example, *M. musculus* possesses two meiosis-specific kleisins, REC8 and RAD21L, and loss of RAD21L (but not REC8) results in a near-complete absence of assembled chromosome axes (Biswas *et al*, 2016). Similarly, in *C. elegans* the REC-8 kleisin is primarily responsible for sister chromosome cohesion, while the kleisin paralogs COH-3 and COH-4 are responsible for chromatin looping and axis assembly (Pasierbek *et al*, 2001; Severson *et al*, 2009; Woglar *et al*, 2020).

While the interactions between HORMADs and axis core complexes are well-understood, a major remaining question is whether and how cohesin complexes interact directly with other chromosome axis proteins. A physical interaction between Red1 and Rec8-containing cohesin complexes has been demonstrated using proximity-labeling and co-immunoprecipation in yeast meiotic cells (Sun *et al*, 2015). Moreover, chromatin immunoprecipitation (ChIP-Seq) experiments have shown that both Red1 and Hop1 strongly co-localize with Rec8-containing cohesin complexes along meiotic chromosomes (Sun *et al*, 2015). Deletion and mutation studies have shown that Red1’s association with meiotic chromosomes depends on both the Hop1 CBR (in nucleosome-dense regions termed “islands”) and on Rec8-containing cohesin complexes (in both islands and in other regions termed “deserts”) (Heldrich *et al*, 2022). Deletion of either *HOP1* or *REC8* partially disrupts Red1’s chromosome association, while a double *HOP1*-*REC8* deletion results in a near-complete loss of Red1 from meiotic chromosomes (Sun *et al*, 2015). Overall, these data support a direct interaction between Red1 and Rec8-containing cohesin complexes, but the molecular basis for this interaction remains unknown.

Here we identify and characterize a direct interaction between *S. cerevisiae* Red1 and Rec8. Using yeast genetics, biochemistry, and AlphaFold 3 structural modeling, we map the Red1-Rec8 binding interface and show that disrupting this interface results in a specific loss of Rec8-dependent Red1 axis association patterns, and compromises the success of meiosis. Based on our findings with *S. cerevisiae* Red1 and Rec8, we identify and validate an equivalent interaction between the *Arabidopsis thaliana* axis core protein ASY3 and the kleisin REC8. Strikingly, we cannot identify an equivalent interaction between mammalian axis core and kleisin proteins, suggesting variability in chromosome axis assembly mechanisms across eukaryotic phyla. Overall, our findings complete the architectural picture of the meiotic chromosome axis in fungi and plants, revealing how cohesin complexes are integrated into the axis to mediate assembly of the compacted chromatin loop array characteristic of meiotic chromosomes.

## Results

### Conserved motifs in Red1’s disordered region are essential for meiosis in *S. cerevisiae*

The *S. cerevisiae* meiotic axis core protein Red1 possesses a folded N-terminal domain with armadillo-repeat (ARM) and pleckstrin homology (PH) subdomains (residues 1-345) (Feng *et al*, 2017), a Hop1-binding closure motif (residues 346-362) (Woltering *et al*, 2000; West *et al*, 2018), and a C-terminal domain that forms coiled-coil homotetramers (residues 733-827; **Figure 1A, Figure S1**) (West *et al*, 2019). Between Red1’s closure motif and coiled-coil region is an extended central region (residues 363-732) that is predicted to be disordered in solution. By analyzing sequence alignments of budding-yeast Red1 proteins, we identified four highly conserved motifs in this region, which we term disordered-region motifs 1-4 (DR1-DR4; **Figure 1B, Figure S2**). To determine whether these motifs are important for meiosis, we replaced the *RED1* gene in *S. cerevisiae* with truncated *red1* constructs lacking each motif (*red1-ΔDR1* through *red1-ΔDR4*) and performed spore viability assays. As previously demonstrated (Rockmill & Roeder, 1988), *S. cerevisiae red1Δ* showed 1% spore viability (n=80) compared to 96% for wild-type *S. cerevisiae* (**Figure 1C**). *RED1* constructs deleted for DR1 or DR2 fully rescued spore viability (to 95% and 93%, respectively), whereas the *red1-ΔDR3* construct showed 15% spore viability, and *red1-ΔDR4* showed 23% spore viability (**Figure 1C**). Thus, *S. cerevisiae* Red1 contains two short motifs in its disordered region, DR3 and DR4, that are important for its function during meiosis.

**Figure 1.**
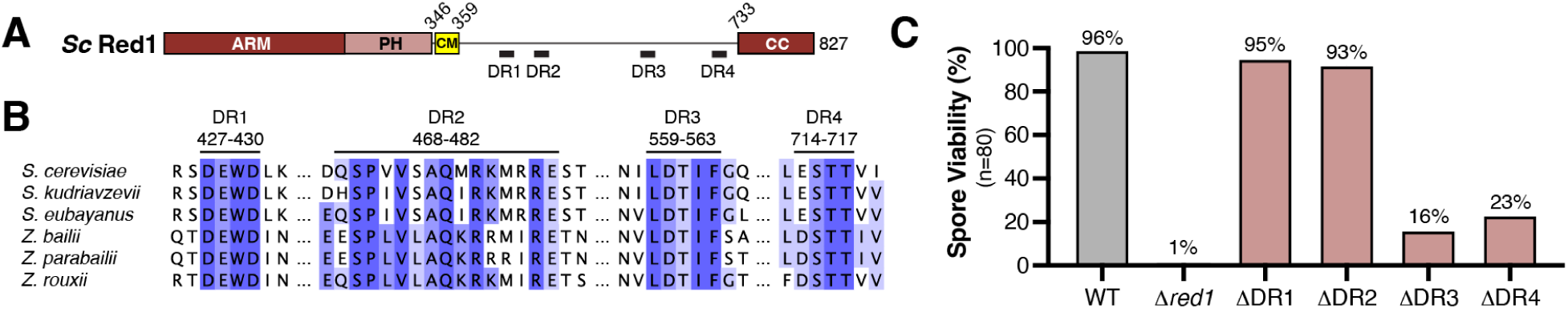
Identification of conserved motifs in the Red1 disordered region. (A) Domain schematic of *S. cerevisiae* Red1. ARM: armadillo repeat domain; PH: pleckstrin homology domain; CM: closure motif; CC: coiled-coil. See **Figure S1** for details. (B) Sequence alignment of conserved motifs DR1-DR4 in the Red1 central disordered region. See **Figure S2** for details and NCBI accession numbers. (C) Spore viability of wild-type *S. cerevisiae* (WT), the *Δred1* deletion mutant, and strains where *RED1* has been replaced with a mutant lacking DR1, DR2, DR3, or DR4. n=80 spores (20 tetrads) per strain.

### Red1 binds the C-terminal winged-helix domain of Rec8

One potential role for Red1 DR3 and/or DR4 is direct binding to another meiotic chromosome-associated protein. To identify potential protein binding partners of Red1 DR3 and DR4, we performed a systematic analysis using AlphaFold 3, which can confidently predict both the structures of individual proteins and interactions between proteins (Abramson *et al*, 2024; Kim *et al*, 2024; Yu *et al*, 2023). We predicted the structure of ∼50 amino acid regions of Red1 containing DR3 or DR4 with a set of 33 meiotic chromosome-associated proteins, including all known subunits of the cohesin complex, chromosome axis and synaptonemal complex components, and proteins responsible for DSB formation and repair (**Figure 2A-B**). We used the interface predicted template modeling (ipTM) score as a metric for the confidence of the predicted interaction, as prior analyses have shown that ipTM correlates well with the accuracy of a predicted protein-protein complex (Genz *et al*, 2025). An ipTM score above 0.8 indicates a highly confident protein complex prediction, while an ipTM score between 0.6 and 0.8 indicates reduced confidence (Abramson *et al*, 2024).

**Figure 2.**
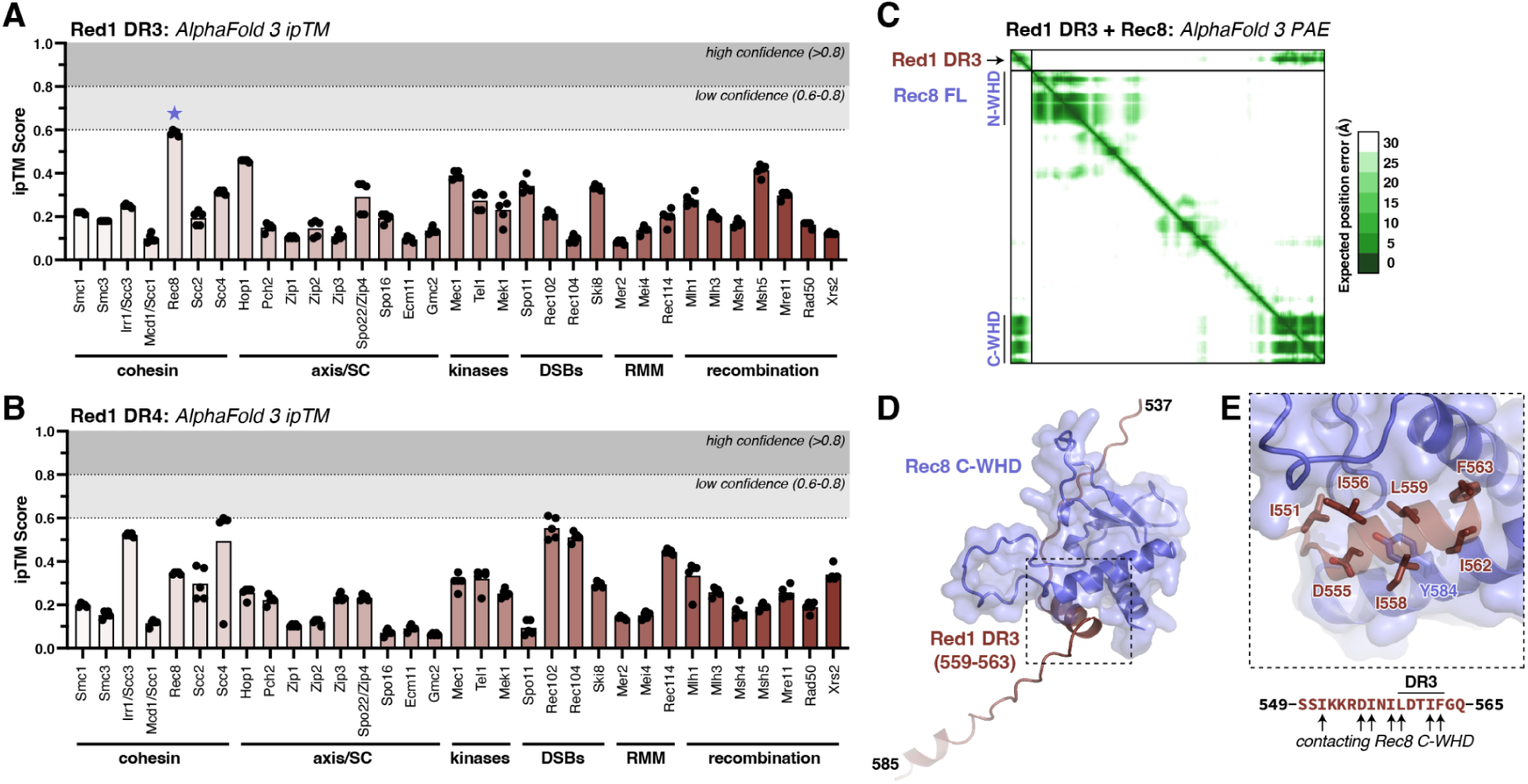
AlphaFold identifies Rec8 as a potential DR3 interaction partner. (A) AlphaFold 3 interface predicted template modeling (ipTM) scores for the *S. cerevisiae* Red1 DR3 region with 33 meiotic chromosome-associated proteins. Black dots indicate ipTM scores for five independent models, and bars indicate average ipTM for all five models. The Red1 DR3-Rec8 pair shown in panels (C)-(E) is marked with a blue star. (B) AlphaFold 3 ipTM scores for the *S. cerevisiae* Red1 DR3 region with 33 meiotic chromosome-associated proteins. The predicted interface with Scc4 was not recapitulated in a prediction of Red1 DR4 with the Scc2-Scc4 complex, and the predicted interface with Rec102 was not recapitulated in a prediction of Red1 DR4 with the Rec102-Rec104 complex (not shown). (C) AlphaFold 3 predicted aligned error (PAE) plot for the Red1 DR3-Rec8 prediction. This plot indicates a predicted interaction between Red1 DR3 and the Rec C-terminal winged helix domain (C-WHD). (D) AlphaFold 3 predicted structure of the Red1 DR3 region (brown) with Rec8 (blue; only C-WHD domain shown). (E) Closeup view of the predicted interface between Red1 DR3 and the Rec8 C-WHD. Residues predicted to bind Rec8 are shown as sticks. *Bottom:* Red1 DR3 region with predicted Rec8-binding residues marked.

The highest-confidence prediction from our AlphaFold interaction screen involved Red1 DR3 and the meiosis-specific cohesin subunit Rec8, with five independent models showing ipTM values between 0.57 and 0.60 (**Figure 2A**). Inspection of the predicted Red1 (DR3)-Rec8 complex showed Red1 DR3 adopting a helical conformation with one hydrophobic face docking against a hydrophobic groove in the Rec8 C-terminal winged helix domain (C-WHD; **Figure 2C-E**). The predicted interface involves three highly conserved residues in DR3 (L559, I562, and F563), plus several less well-conserved residues upstream of DR3 (I551, D555, I556, and I558; **Figure 2E**). The predicted Red1-binding interface on the Rec8 C-WHD is highly conserved and predominantly hydrophobic (**Figure S3A-C**), and is distinct from the equivalent surface on the mitotic kleisin Mcd1 (Scc1; **Figure S3D**). AlphaFold predictions of the equivalent complex from *S. pombe* (Rec10 and Rec8) show a similar interface between a short amphipathic α-helix in Rec10’s central disordered region and the Rec8 C-WHD (**Figure S4**).

We next sought to directly test binding between Red1 DR3 and the Rec8 C-WHD (residues 574-680). Since the Rec8 C-WHD was unstable when purified on its own (not shown), we reconstituted a stable complex of the Rec8 C-WHD bound to the Smc1 ATPase head domain (residues 1-214 and 1024-1225 fused with an eight-residue linker; see **Methods**) (**Figure 3A, Figure S5**). We separately purified GST-tagged Red1 DR3 (residues 546-569), and performed an affinity pulldown with glutathione resin. In this assay, wild-type Red1 DR3 bound the Smc1(head)+Rec8(C-WHD) complex, but a mutant with I558 and L559 each mutated to lysine (I558K/L559K; termed Red1 DR3-KK) failed to bind (**Figure 3B**). We also generated a point mutant of Rec8 in the Red1 binding interface (Y584D), and found that this mutant disrupted Red1 DR3 binding (**Figure 3B**). Finally, we found that excess Red1 DR3 peptide competed for Rec8(C-WHD)-Smc1(head) binding, but that a Red1 DR3-KK mutant peptide did not compete for binding (**Figure 3B**). These data show that Red1 DR3 directly interacts with the Rec8(C-WHD) as predicted by AlphaFold.

**Figure 3.**
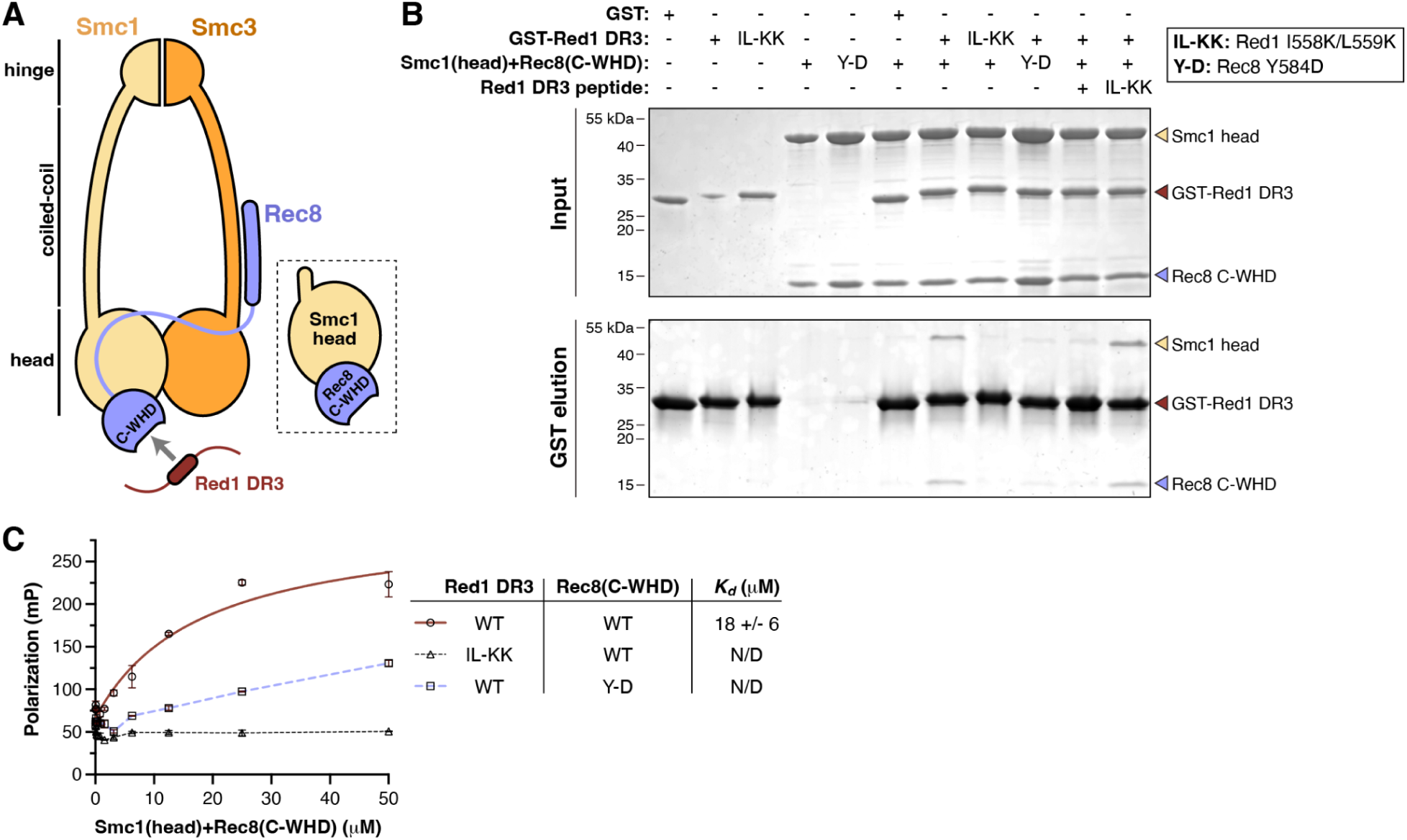
Red1 DR3 binds Rec8. (A) Schematic of an Smc1-Smc3-Rec8 cohesin complex binding Red1 DR3, based on an AlphaFold 3 prediction of the full complex (**Figure S5**). *Inset:* Schematic of Smc1(head)+Rec8(C-WHD) subcomplex used for affinity pulldown assays. (B) Affinity pulldown assay showing interaction of GST-Red1 DR3 with the Smc1(head)+Rec8(C-WHD) subcomplex. Top gel: 10% inputs. Bottom gel: glutathione affinity resin elution. Both gels are visualized by Coomassie blue staining. IL-KK: Red1 I558K/L559K. Y-D: Rec8 Y584D. (C) Fluorescence polarization binding assay between the Smc1(head)+Rec8(C-WHD) complex (WT or Rec8 Y584D) and fluorescein-labeled Red1 DR3 peptides (WT or IL-KK mutant).

To complement the above result, we performed fluorescence polarization binding assays using the reconstituted Smc1(head)+Rec8(C-WHD) complex and fluorescein-labeled Red1 DR3 peptides. We found that wild-type Red1 DR3 bound the cohesin subcomplex with a dissociation constant (*K_d_*) of ∼18 μM, and a Red1 DR3-KK mutant peptide did not show detectable binding (**Figure 3C**). As in our pulldown assays, we found that the Rec8 Y584D mutation eliminated binding to the Red1 DR3 peptide (**Figure 3C**). Finally, we tested spore viability for a diploid yeast strain with both copies of *RED1* replaced with a *red1-KK* (I558K/L559K) mutant, and found that spore viability was 46% (n=80). This intermediate spore viability is consistent with a partial loss of binding between Red1 and Rec8. Based on these findings and consistent with a contemporary report from Murakami and colleagues (Rajalingam *et al*, 2026), we term DR3 the Red1 cohesin-interacting motif (CIM).

### The Red1 CIM mediates Rec8-dependent axis association

To test the role of the Red1 CIM in chromosome axis assembly, we performed chromosome spreads of *S. cerevisiae* cells carrying the *red1-KK* mutation in pachytene (four hours after induction of meiosis). Compared to wild-type cells, *red1-KK* cells showed reduced levels of chromosome-associated Red1 as judged by immunostaining (**Figure 4A-B**), but retained robust synaptonemal complex assembly as judged by localization of the central element protein Gmc2 (**Figure 4A,C**). Western blot analysis indicated equivalent total Red1 levels in wild-type and *red1-KK* cells (**Figure 4D**), indicating that the reduction in chromosomal Red1 is not a result of reduced protein expression or stability.

**Figure 4.**
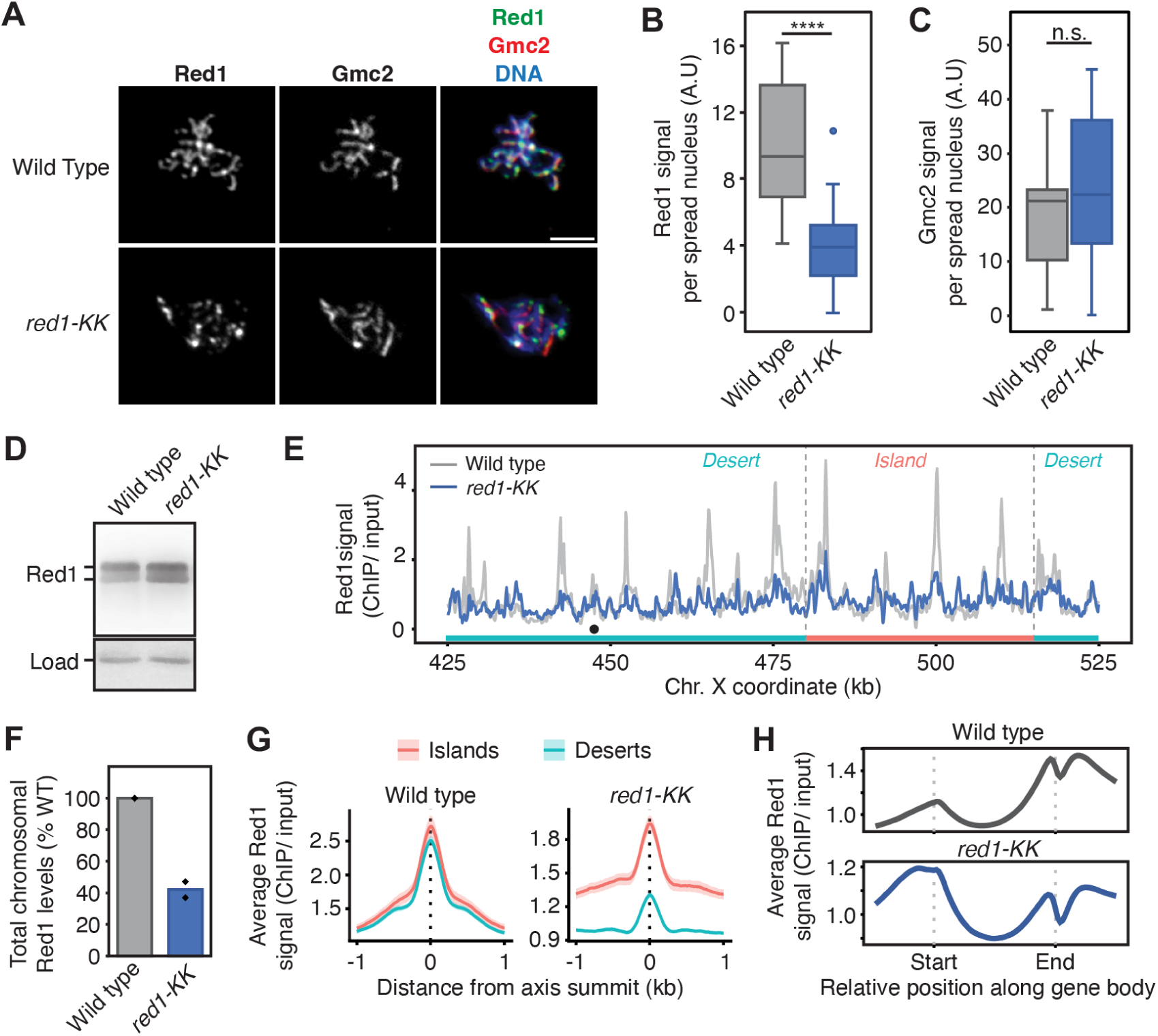
The *red1-kk* mutant disrupts Red1 chromosome localization and Rec8-dependent axis association. (A) Chromosome spreads showing localization of Red1 and the synaptonemal complex component Gmc2, in wild type and *red1-kk* (I558K/L559K) mutant strains. Scale bar = 0.5 µm. (B) Quantification of chromosome-associated Red1 fluorescence signal from chromosome spreads (n = 15 and 20 cells, respectively). Statistical significance was calculated using Welch’s t-test (**** indicates p = 2.7*10^-5^). (C) Quantification of chromosome-associated Gmc2 fluorescence signal from chromosome spreads (n = 16 and 22 cells, respectively). Statistical significance was calculated using Welch’s t-test (n.s. indicates not significant; p = 0.38). (D) Western blot showing overall Red1 levels in wild type and *red1-kk* strains. Loading control: cross-reacting band of a histone H2A western blot. (E) Red1 ChIP-Seq for wild type (gray) and *red1-KK* (blue) in a region of chromosome X with an island region (salmon) and two desert regions (cyan). Black dot indicates the location of the centromere. (F) Total chromosomal Red1 in *red1-KK* (blue) compared to wild type (grey), from spike-in normalized ChIP-Seq (SNP-ChIP). (G) Average Red1 enrichment at axis attachment sites (Sun *et al*, 2015) in wild type (left) and *red1-KK* (right) separated into islands (salmon) and deserts (cyan). The 95% confidence interval (C.I.) for the average lines is shown. (H) Metagene analysis of Red1 in wild type (top; gray) and *red1-KK* (bottom; blue) indicating average ChIP-seq enrichment along genes. The 95% C.I. for the average lines is shown.

We next performed ChIP-seq analysis to assess Red1-chromosome association patterns. In wild-type cells, Red1 localization depends on two factors: Rec8-containing cohesin complexes and the Hop1 CBR (Heldrich *et al*, 2022; Milano *et al*, 2024). In nucleosome-dense regions termed “islands”, the Hop1 CBR promotes association of both Hop1 and Red1 to chromatin.

Layered onto that pattern, Rec8 promotes peaks of Red1 (and Hop1) localization at cohesin-bound sites along the entire genome, including islands and less nucleosome-rich regions termed “deserts” (Heldrich *et al*, 2022; Milano *et al*, 2024). In the *red1-KK* mutant, Red1 enrichment at Rec8-dependent peaks was strongly reduced (**Figure 4E**), resulting in an overall reduction of chromosome-associated Red1, as measured by spike-in controlled ChIP (**Figure 4F**). Red1 binding was more severely reduced in deserts compared with islands (**Figure 4E,G**), consistent with a specific loss of Rec8-dependent Red1 axis association. *red1-KK* cells also showed less pronounced enrichment of Red1 at the 3ʹ ends of genes (**Figure 4H**), a pattern that is also observed with Rec8 and has been attributed to active RNA polymerase shifting cohesin complexes toward the end of transcripts (Heldrich *et al*, 2022). Overall, these data support a model in which the *red1-KK* mutant causes a specific loss of Rec8-dependent Red1 localization to chromosomes, while maintaining Hop1-dependent localization. However, we note that Red1 binding in the *red1-kk* mutants differs significantly from *rec8Δ* mutants, in which Red1 localization in desert regions is almost completely lost (Sun *et al*, 2015; Heldrich *et al*, 2022).

We hypothesize that this difference may arise from two sources: First, even though we observe a loss of binding *in vitro* with the *red1-KK* mutant, this mutant may retain residual Red1-Rec8 interaction in cells. Second, the presence of functional Rec8 in *red1-KK* cells means that overall chromosome architecture as a linear array of chromatin loops is maintained, compared to a complete loss of cohesin function in *rec8Δ* cells. This organization may have an indirect effect on the assembly of Red1 and Hop1 on chromosomes.

### Plant ASY3 contains a REC8-binding CIM

We previously showed that meiotic axis core proteins from both plants (*Arabidopsis thaliana* ASY3) and mice (*Mus musculus* SYCP2) possess HORMAD-binding closure motifs near their N-termini and C-terminal coiled-coil domains that form 2:2 heterotetramers with small coiled-coil protein partners (*A. thaliana* ASY4 and *M. musculus* SYCP3, respectively) (West *et al*, 2019).

We sought to determine if *A. thaliana* ASY3 and *M. musculus* SYCP2 bind meiotic cohesin complexes in a Red1-like manner. Using yeast two-hybrid assays, we tested for binding of *A. thaliana* ASY3 (full-length and truncations) to either the *A. thaliana* SMC1 head domain or a complex of the SMC1 head plus the C-WHD domain of *A. thaliana* REC8 (also termed SYN1) (Bai *et al*, 1999; Bhatt *et al*, 1999; Cai *et al*, 2003). We detected binding between full-length ASY3 and the SMC1(head)+REC8(C-WHD) complex but not the SMC1(head) alone, indicating that ASY3 likely binds the REC8 C-WHD domain (**Figure 5A-B, Figure S6A-C**). Using a series of ASY3 truncations, we narrowed the likely REC8-binding region to residues 180-230 (**Figure 5A-C, Figure S6A-D**). Using a multiple sequence alignment of plant ASY3 homologs, we identified a conserved motif spanning roughly residues 196-204 of *A. thaliana* ASY3, which contains several highly conserved hydrophobic residues (**Figure S6E**). We generated a quadruple mutant with four hydrophobic residues mutated to polar or negatively charged amino acids (LLIL mutant: L196D/L200D/I203N/L204E), and found that this mutant eliminated the interaction between ASY3 and the SMC1(head)+REC8(C-WHD) complex (**Figure 5C, Figure S6D**). These data show that, like *S. cerevisiae* Red1, *A. thaliana* ASY3 possesses a CIM in its central disordered region. We performed parallel yeast two-hybrid assays with *M. musculus* SYCP2 and the C-WHD domains of both REC8 and RAD21L, but did not detect an interaction (**Figure S7**).

**Figure 5.**
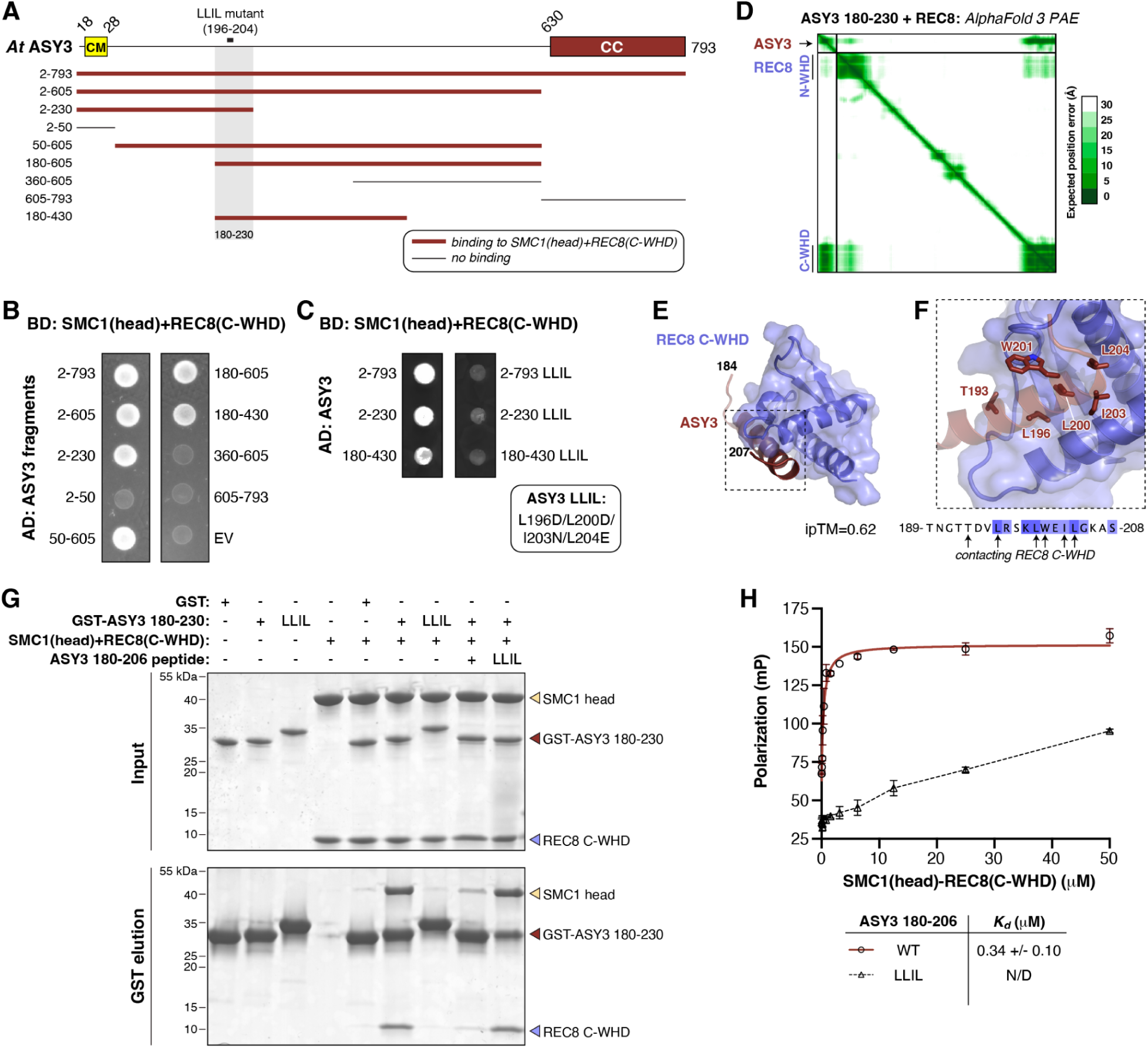
The axis core-kleisin interface is conserved in plants. (A) Domain schematic of *Arabidopsis thaliana* ASY3, with N-terminal closure motif (CM) and C-terminal coiled-coil (CC) domains noted (West *et al*, 2019). Bottom: results of yeast two-hybrid screens with noted ASY3 fragments and the SMC1(head)+REC8(C-WHD) complex (see panel (B)), with thick brown lines indicating a positive interaction and thin black lines indicating a negative interaction. The inferred minimal binding region (ASY3 180-230) is shaded, with the location of the ASY3 LLIL mutant (L196D/L200D/I203N/L204E) noted. See **Figure S6A-C** for full results including ASY3 fragment interactions with SMC1(head) alone, REC8(C-WHD) alone, and the SMC1(head)+REC8(C-WHD) complex. (B) Yeast two-hybrid data for ASY3 fragments and the SMC1(head)+REC8(C-WHD) complex. AD: activation domain fusion; BD: DNA binding domain fusion; EV: empty vector. Growth on selection plates (-LTH: lacking leucine, tryptophan, and histidine) is shown; see **Figure S6** for growth on non-selective and stringently selective plates. (C) Yeast two-hybrid data for ASY3 fragments and their LLIL mutants, and the SMC1(head)+REC8(C-WHD) complex. See **Figure S6D** for full results. (D) AlphaFold 3 predicted aligned error (PAE) plot for the ASY3 180-230-REC8 prediction. This plot indicates a predicted interaction between ASY3 ∼190-200 and the REC8 C-terminal winged helix domain (C-WHD). (E) AlphaFold 3 predicted structure of ASY3 residues 184-207 (brown) with REC8 (blue; only C-WHD domain shown). (F) Closeup view of the predicted interface between ASY3 and the REC8 C-WHD. Residues predicted to bind REC8 are shown as sticks. *Bottom:* ASY3 residues 189-208 (colored by conservation in an alignment of plant ASY3 proteins; see **Figure S6E**) with predicted REC8-binding residues marked. (G) Affinity pulldown assay showing interaction of GST-ASY3 180-230 with the SMC1(head)+REC8(C-WHD) subcomplex. Top gel: 10% inputs. Bottom gel: glutathione affinity resin elution. Both gels are visualized by Coomassie blue staining. (H) Fluorescence polarization binding assay between the SMC1(head)+REC8(C-WHD) complex and fluorescein-labeled ASY3 180-206 peptides (WT or LLIL mutant).

To gain further information on the *A. thaliana* ASY3-REC8 interaction, we performed an AlphaFold 3 structure prediction with *A. thaliana* ASY3 180-230 and full-length REC8 (**Figure 5D-F**). The resulting prediction showed a low-confidence interaction (ipTM = 0.62), with the conserved motif of ASY3 we identified adopting an ɑ-helical structure and binding the REC8 C-WHD equivalently to our predictions of the *S. cerevisiae* Red1-Rec8 interaction (**Figure 5E**). All four residues mutated in the LLIL mutant (L196, L200, I203, and L204) are docked against a hydrophobic surface of REC8 in the predicted structure (**Figure 5F**). Parallel AlphaFold 3 predictions of *M. musculus* SYCP2 with REC8 or RAD21L showed no interaction (not shown), suggesting that the detected axis core-kleisin interaction is not conserved in the mouse.

We next reconstituted the *A. thaliana* SMC1(head)+REC8(C-WHD) complex, and performed affinity pulldown binding assays with GST-tagged ASY3 180-230. We found that ASY3 180-230, but not the LLIL mutant, robustly interacts with the SMC1(head)-REC8(C-WHD) complex (**Figure 5G**). This interaction was sensitive to incubation with a competitor ASY3 peptide spanning residues 180-206, but not an LLIL mutant peptide (**Figure 5G**). Finally, we used fluorescence polarization to directly measure the affinity of the interaction between ASY3 and the SMC1(head)+REC8(C-WHD) complex. We detected binding with an affinity (*K_d_*) of 0.3 µM for the wild-type ASY3 180-206 peptide, and no interaction using an LLIL mutant peptide (**Figure 5H**). Overall, these data show that plant ASY3 binds the meiotic cohesin complex equivalently to budding yeast Red1.

## Discussion

A central question in meiosis is how chromosomes are organized by the chromosome axis, and how DNA cleavage and recombination are controlled by axis proteins to give rise to interhomolog crossovers. To answer this question, it is essential to dissect the detailed molecular interactions between proteins of the chromosome axis, their interactions with DNA cleavage and recombination factors, and their later interactions with proteins that make up the central region/transverse filaments of the synaptonemal complex. Here, we identify a specific interaction between the filamentous axis core proteins and the meiosis-specific cohesin subunit Rec8, that is conserved across fungi and plants. In the budding yeast *S. cerevisiae*, we find that disrupting the Red1-Rec8 interface results in a specific loss of Rec8-dependent Red1-chromosome association in desert regions of chromosomes, while maintaining Hop1-dependent localization in island regions.

When combined with prior work defining how Red1 oligomerizes and binds the HORMAD protein Hop1, and how both cohesin and Hop1 bind DNA, we can assemble a molecular model of meiotic chromosome axis architecture in *S. cerevisiae* (**Figure 6**). We propose that the axis core protein Red1 forms filaments through its C-terminal coiled-coil region that contain a high concentration of cohesin binding sites. Cohesin complexes are captured into this structure by interacting with the Red1 CIM, thereby stabilizing chromosomes into a linear array of chromatin loops. It is possible that Red1 binding increases the lifetime of cohesin-DNA binding and/or alters its loop extrusion dynamics to further stabilize this structure. To perform this action, Red1 needs to interact with Hop1 through its closure motif, and these proteins together recruit and control the activity of Spo11 complexes to generate DSBs, and recombination proteins to repair DSBs as interhomolog crossovers. Assembly of the synaptonemal complex, mediated by as-yet unknown interactions between axis proteins and transverse filament proteins (in *S. cerevisiae*, Zip1), recruits the AAA+ ATPase Pch2 to remove most Hop1 from the axis, thereby shutting down DSB formation and allowing repair of remaining breaks via the sister chromosome. Major remaining questions with regard to axis architecture include: (1) which population of cohesin complexes – dynamic loop-extruding complexes, cohesive complexes, or both – interacts with Red1; and (2) what is the role of nucleosomal DNA binding by Hop1’s central chromatin binding domain (CBR)?

**Figure 6.**
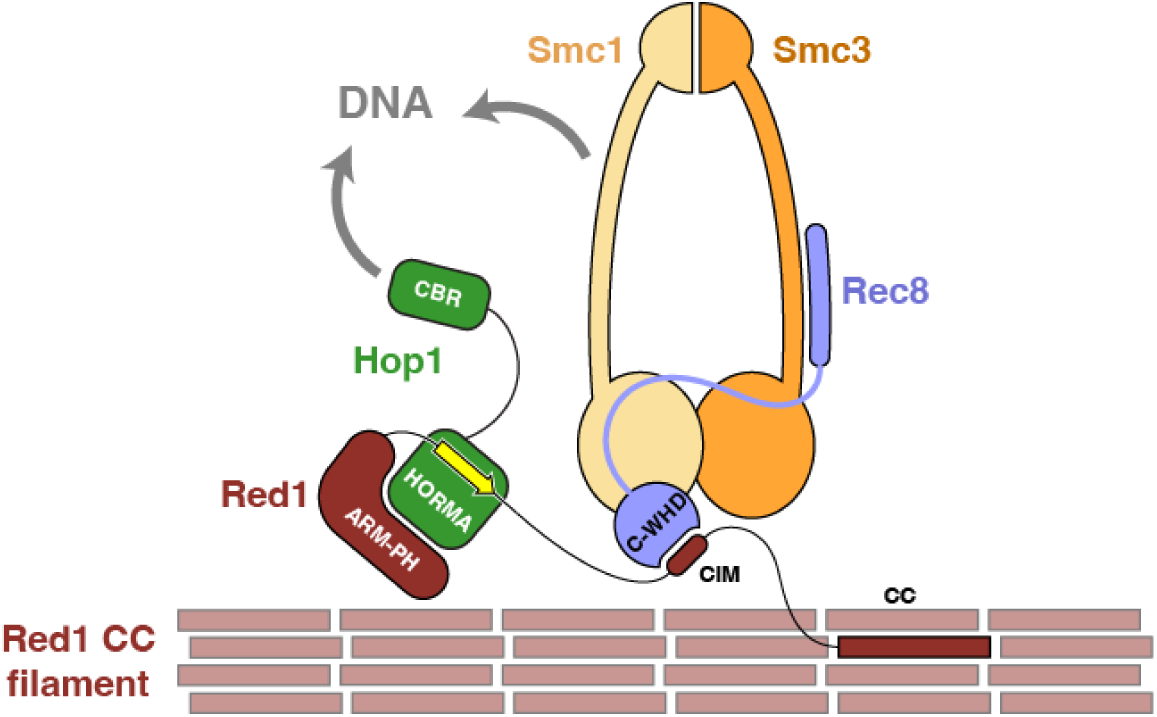
Molecular architecture of the *S. cerevisiae* chromosome axis. Schematic of the *S. cerevisiae* chromosome axis, with Red1 brown, Hop1 green, and cohesin complexes yellow (Smc1), orange (Smc3), and blue (Rec8; C-terminal winged helix domain (C-WHD) labeled). Other subunits of cohesin are not shown. The Red1 ARM-PH, CIM, and coiled-coil (CC) regions are labeled, and the closure motif is shown in yellow. The Hop1 HORMA domain and chromatin binding region (CBR) are labeled. Both cohesin complexes and the Hop1 CBR interact with chromosomal DNA.

The axis core protein-cohesin interface we identified is conserved across fungi and plants, but likely not in mammals. Thus, a distinct axis-cohesin interaction must underlie axis architecture in mammals. Moreover, some organisms including the nematode *C. elegans* entirely lack axis core proteins homologous to fungal Red1, plant ASY3, and mammalian SYCP2. We propose that in *C. elegans*, the synaptonemal complex - which assembles prior to DSB formation in this organism - functionally replaces axis core proteins’ role in capturing and organizing cohesin complexes to generate linear arrays of chromatin loops. Still unknown, however, is how *C. elegans* meiotic HORMAD proteins are recruited to the chromosome axis in the absence of axis core proteins. Thus, more work is required to comprehensively define axis architecture in diverse eukaryotic phyla.

In addition to the CIM, our *red1* deletion screen identified another motif (DR4; residues 714-717) as important for successful meiosis, with deletion of this motif resulting in 23% spore viability). The sequence of DR4 (ESTT) suggests that this motif may be phosphorylated. We speculate that Red1 DR4 phosphorylation may regulate protein-protein interactions with an as-yet unidentified partner protein. Alternatively, given the proximity of DR4 to Red1’s C-terminal coiled coil domain, DR4 phosphorylation could modulate Red1 filament formation, perhaps enabling axis remodeling specifically at recombination sites or mediating axis disassembly in late meiotic prophase. Further work will be required to define the role of the Red1 DR4 motif, and whether, like the CIM, its role is conserved in a wider set of organisms.

## Materials and Methods

### Yeast strain construction

All *S. cerevisiae* strains generated in this study are listed in **Table 1**. All yeast strains for spore viability assays are in the *S. cerevisiae* SK1 background *(ho::LYS2, lys2, ura3, leu2:hisG, his3::hisG, trp1::hisGSK1, GAL1-).* Unless otherwise noted, yeast cells were cultured in YPD media (1% w/v yeast extract, 2% w/v peptone, 2% w/v dextrose). Gene deletions and integration of *RED1* (wild type and mutant) linear plasmids into the *ura3* locus in SK1 strains for spore viability assays were performed using standard PCR-based homologous recombination (Janke *et al*, 2004; Longtine *et al*, 1998).

**Table 1.** Yeast strains used in this study.

| Strain | Genotype | Ploidy |
| --- | --- | --- |
| KDC306 | *Mata*, *trp1-901*, *leu2-3*, *112*, *ura3-5 2*, *his3-200*, *gal4delta*, *gal80delta*, *LYS2::GAL1(UAS)-GAL1(TATA)-HIS3*, *GAL2(UAS)-GAL2(TATA)-ADE2*, *URA3::MEL1(UAS)-MEL1(TATA)-lacZ*, *MEL1* | Haploid |
| KDC307 | *Mata<sub>α</sub>*, *ura3-52*, *his3-200*, *ade2-101*, *trp1-901*, *leu2-3*, *112*, *gal4delta*, *met-*, *gal80delta*, *MEL1*, *URA3::GAL1(UAS)-GAL1(TATA)-lacZ* | Haploid |
| KDC833 | *red1Δ::KanMx*, empty vector::*URA*, *Mata/Mata<sub>α</sub>*, *ho::LYS2*, *lys2*, *ura3/URA*, *leu2::hisG:LEU2*, *his3::hisG*, *trp1::hisGSK1:TRP1/ wild-type*Note: *GAL1-*. | Diploid |
| KDC834 | *red1Δ::KanMx*, *RED1 WT::URA*, *Mata/Mata<sub>α</sub>*, *ho::LYS2*, *lys2*, *ura3/URA*, *leu2::hisG:LEU2*, *his3::hisG*, *trp1::hisGSK1:TRP1/ wild-type*Note: *GAL1-*. | Diploid |
| KDC837 | *red1Δ::KanMx*, *red1 DR1Δ (aa 427-430 deleted)::URA*, *Mata/Mata<sub>α</sub>*, *ho::LYS2*, *lys2*, *ura3/URA*, *leu2::hisG:LEU2*, *his3::hisG*, *trp1::hisGSK1:TRP1/ wild-type*Note: *GAL1-*. | Diploid |
| KDC838 | *red1Δ::KanMx*, *red1 DR2Δ (aa 468-482 deleted)::URA*, *Mata/Mata<sub>α</sub>*, *ho::LYS2*, *lys2*, *ura3/URA*, *leu2::hisG:LEU2*, *his3::hisG*, *trp1::hisGSK1:TRP1/ wild-type*Note: *GAL1-*. | Diploid |
| KDC839 | *red1Δ::KanMx*, *red1 DR3Δ (aa 559-563 deleted)::URA*, *Mata/Mata<sub>α</sub>*, *ho::LYS2*, *lys2*, *ura3/URA*, *leu2::hisG:LEU2*, *his3::hisG*, *trp1::hisGSK1:TRP1/ wild-type*Note: *GAL1-*. | Diploid |
| KDC840 | *red1Δ::KanMx*, *red1 DR4Δ (aa 714-717 deleted)::URA*, *Mata/Mata<sub>α</sub>*, *ho::LYS2*, *lys2*, *ura3/URA*, *leu2::hisG:LEU2*, *his3::hisG*, *trp1::hisGSK1:TRP1/ wild-type*Note: *GAL1-*. | Diploid |
| KDC877 | *red1Δ::KanMx*, *red1<sup>I558K, L559K</sup>::URA*, *Mata/Mata<sub>α</sub>*, *ho::LYS2*, *lys2*, *ura3/URA*, *leu2::hisG:LEU2*, *his3::hisG*, *trp1::hisGSK1:TRP1/ wild-type*Note: *GAL1-*. | Diploid |

URA-integrating yeast plasmids for introducing wild type or mutant *RED1* genes into their respective *red1Δ* SK1 strains were constructed by PCR-amplifying the full-length wild type gene (*RED1)* gene along with upstream (5’) promoter sequence (700 bp) from *S. cerevisiae* S288C genomic DNA and cloning it into the *SmaI* site of the YIplac211 integrating vector (a gift from the Kellogg Lab, UC Santa Cruz). Internal deletions of *RED1* were introduced via PCR and isothermal assembly, and point mutations in both genes were introduced using PCR-based mutagenesis.

To generate diploid strains for spore viability assays, *red1Δ* mutant SK1 strains were marked with selectable auxotrophies: *MATα* strains were made *TRP^+^* and *MATa* strains were made *LEU^+^* via PCR-based homologous recombination at the *leu2* locus. Wild type and mutant *RED1* carried on linearized YIplac211 plasmids (digested with StuI) were then integrated at the *ura3* locus of the appropriate *MATa* or *MATα* haploids. Transformants were selected on CSM-URA plates. Correct integration was confirmed by PCR and by verifying the presence of *RED1* coding sequences from phenol-chloroform extracted genomic DNA. Confirmed haploid strains were mated on YPD plates, and diploids were selected on CSM-LEU-TRP plates.

### Spore viability assays

Diploid strains were grown to log phase in YPD media (OD_600_ 0.4-0.6) at 30°C. For each strain, a 1 mL saturated culture in YPD was pelleted by centrifugation, then resuspended in 5 mL of sporulation media (1% potassium acetate solution) and incubated at 30°C for 3-5 days with shaking at 250 rpm. After sporulation, 300 μL of cells were pelleted and resuspended in 1 M sorbitol containing zymolyase (1 mg/mL final concentration) and incubated at room temperature for 5 minutes. Treated cells were then diluted with water, and 15 μL of suspension was applied to the edge of a YPD agar plate. The plate was tilted to allow excess liquid to drain and incubated at 30°C for 15 minutes to allow the sample to soak in. Twenty tetrads were dissected and plated on the same YPD plate. Plates were incubated at 30°C for 2-3 days, after which spore viability was determined as the number of viable spores divided by the total number of spores plated (n=80).

### Synchronous meiosis

To induce synchronous meiosis, cells were grown in liquid YPD media for 24 hours at room temperature. Saturated cultures were diluted to O.D._600_ = 0.3 in BYTA media (50 mM potassium phthalate, 1% yeast extract, 2% tryptone, 3% potassium acetate) and grown for 16.5 hours at 30°C. Cells were washed twice with water and resuspended in SPO media (0.3% potassium acetate) and incubated at 30°C to start the meiotic time course (t = 0 hours). Synchrony of meiotic entry was confirmed by analyzing DNA content changes over time using flow cytometry.

### AlphaFold 3 interaction screening

For identification of potential interactions using AlphaFold 3, the structures of *S. cerevisiae* Red1 DR3 or DR4 motifs (plus 20 additional residues at the N- and C-termini of each motif) were predicted with each potential binding partner at 1:1 stoichiometry using the AlphaFold 3 web server (https://alphafoldserver.com) (Abramson *et al*, 2024). All results were downloaded and the ipTM scores for each of five models were extracted from the [jobname]_summary_confidences_[modelnumber].json file. ipTM scores for each model were plotted and the protein pairs with the highest ipTM scores were manually inspected.

To identify potential interactions between meiotic axis core proteins and cohesin subunits, complex protein models were generated using the AlphaFold3 public server with multimer mode enabled. For *S. cerevisiae*, the DR3 region of Red1 (residues 546-569) was modeled against full-length sequences of meiotic chromosome-associated proteins, including all four cohesin subunits (Smc1, Smc3, Rec8, and Scc3). Protein sequences were retrieved from the UniProt sequence database and submitted in FASTA format, with each pair modeled as a heterodimer using default parameters. For *A. thaliana*, the full-length ASY3 protein was divided into fragments-based on guidance from West et al. (2019)- to ensure full sequence coverage. Each fragment was individually screened against the C-terminal winged-helix domain of the meiotic kleisin subunit REC8 (residues 551-627) using the same AlphaFold3 settings. For each model protein pair, the predicted ipTM score was used to determine the likelihood of a direct interaction. Scores >0.6 were considered as potential direct and real protein interactions. Models were ranked by ipTM scores, and select axis core-cohesin complex models were visualized in PyMOL to examine binding interfaces and guide site-directed mutagenesis of Red1/ASY3 and Rec8/REC8 for downstream spore viability, biochemical, and Y2H assays.

### Cloning, expression, and purification of axis core and cohesin proteins

To reconstitute the *S. cerevisiae* Smc1(head)+Rec8(C-WHD) complex, a co-expression vector was created using UC Berkeley Macrolab vector 2B-T (Addgene #29666) encoding the Rec8 C-WHD (residues 574-680) with a TEV protease-cleavable N-terminal His_6_-tag, plus untagged Smc1(head) (residues 1-214 and 1024-1225, fused with a GSGSASAG linker). Separately, Red1 DR3 (residues 546-569) was cloned into UC Berkeley Macrolab vector 2G-T (Addgene #29707) to generate a TEV protease-cleavable N-terminal His_6_-GST (glutathione S-transferase) fusion. Point mutations were generated by PCR mutagenesis.

To reconstitute the *A. thaliana* SMC1(head)+REC8(C-WHD) complex, a co-expression vector was created using UC Berkeley Macrolab vector 2B-T encoding the REC8 C-WHD (residues 551-627) with a TEV protease-cleavable N-terminal His_6_-tag, plus untagged SMC1(head) (residues 1-181 and 1023-1218, fused with a GSGSASAG linker). Separately, an ASY3 truncation encoding residues 180-230 was cloned into UC Berkeley Macrolab vector 2G-T (Addgene #29707) to generate a TEV protease-cleavable N-terminal His_6_-GST (glutathione S-transferase) fusion. Point mutations were generated by PCR mutagenesis.

For expression of GST-tagged Red1 or ASY3 fragments, plasmids were transformed into *E. coli* Rosetta2 (DE3) pLysS cells (EMD Millipore) and grown overnight in liquid LB media plus chloramphenicol (35 µg/mL) and carbenicillin (100 µg/mL) at 37°C with shaking. 5 mL saturated overnight culture was used to inoculate each of 2 x 1L 2XYT media plus chloramphenicol and carbenicillin in 2L Erlenmeyer flasks and grown at 37°C with sharking at 180 RPM until the OD_600_ reached 0.6-0.8 (4-6 hours). Cultures were cooled on ice for 10 minutes and protein expression was induced by adding IPTG (isopropyl β-D-1-thiogalactopyranoside) at 0.6 mM, then incubated a further 16 hours at 20°C. Cells were harvested by centrifugation, resuspended in buffer A (20 mM Tris-HCl pH 7.5, 300 mM NaCl, 2 mM β-mercapethanol, and 10% glycerol) plus 10 mM imidazole pH 8 and protease inhibitors (PMSF, leupeptin, pepstatin, and aprotinin), and lysed by sonication (Branson Sonifier). Lysates were clarified by centrifugation, and the supernatant was applied to a 2 mL bed-volume Ni^2+^ affinity column (Ni-NTA Superflow, Qiagen). The column washed with 10 column volumes of buffer A plus 20 mM imidazole pH 8, then 10 column volumes of buffer B (20 mM Tris-HCl pH 7.5, 150 mM NaCl, 2 mM β-mercapethanol, and 10% glycerol) plus 20 mM imidazole pH 8, then eluted with 5 column volumes of buffer B plus 300 mM imidazole pH 8. Purified protein was concentrated (Amicon Ultra, EMD Millipore) then applied to a size exclusion column (Superdex 200 Increase 10/300, Cytiva) in buffer GF (50 mM Tris-HCl pH 8, 150 mM NaCl, 10% glycerol, and 1 mM DTT). Fractions containing the protein of interest were pooled, concentrated, aliquotted, and stored at -80°C for biochemical assays.

For expression of *S. cerevisiae* Smc1(head)+Rec8(C-WHD) and *A. thaliana* SMC1(head)+REC8(C-WHD) complex, the same procedure was followed as above with the following differences: Proteins were expressed in LOBSTR (DE3) RIL cells (Kerafast) (Andersen *et al*, 2013), 4 x 1L 2XYT cultures were used for protein expression, and proteins were further purified by anion exchange chromatography (5 mL HiTrap Q HP, Cytiva) after Ni2+ affinity and before size exclusion chromatography, using a gradient from buffer B (20 mM Tris-HCl pH 7.5, 150 mM NaCl, 2 mM β-mercapethanol, and 10% glycerol) to buffer B supplemented with an additional 850 mM NaCl (for a total of 1M NaCl).

### Glutathione pulldown assays

For pulldown assays, 25 µg wild type or mutant GST-tagged axis core proteins (*S. cerevisiae* Red1 or *A. thaliana* ASY3) were incubated with 35 µg of their cognate cohesin subcomplexes (25 µL total reaction volume) in pulldown buffer (50 mM Tris-HCl pH 7.5, 150 mM KCl, and 10% glycerol) for 60 minutes at 4°C. For assays that include competitor peptides, peptides were added at 10-fold molar excess relative to the cohesin sub-complexes. 30 µL of 50% glutathione bead slurry (New England Biolabs) in pulldown buffer was added, gently mixed, and incubated a further 1 hour at 4°C with gentle rotation. Beads were pelleted by centrifugation, then washed three times with 1 mL of wash buffer (50 mM Tris-HCl pH 7.5, 150 mM KCl, and 10% glycerol) with pelleting between each wash. Bound complexes were eluted in 15 µL of elution buffer (50 mM Tris-HCl pH 7.5, 150 mM KCl, 10 mM reduced L-glutathione, and 10% glycerol).

For analysis, 5% (5 µL) input samples were collected prior to addition of glutathione bead slurry, and 100% (12 µL) elution samples were collected from elutions. Both input and eluted samples were mixed with 6x SDS sample buffer (420 mM Tris-HCl,18% SDS, 60% glycerol, and 30% β -mercapethanol), boiled for 5 minutes at 95°C, and separated on a 10% polyacrylamide gel. Gels were stained with Coomassie Brilliant Blue to visualize protein bands.

### Fluorescence polarization assays

Synthesized peptides were obtained from Biomatik with an N-terminal 6-FAM (fluorescein) fluorophore. Peptide sequences:

Red1 DR3 (residues 546-569): KSNSSIKKRDINILDTIFGQPPSK

Red1DR3 mutant (I558K/L559K): KSNSSIKKRDIN<u>KK</u>DTIFGQPPSK

ASY3 (residues 180-206): EKMDKPGKETNGTTDVLRSKLWEILGK

ASY3 LLIL mutant: EKMDKPGKETNGTTDV<u>D</u>RSK<u>D</u>WE<u>NE</u>GK

Binding reactions (20 μl) contained 20 nM peptide and varying concentrations of cohesin subcomplexes in binding buffer (50 mM Tris-HCl pH 7.5, 150 mM NaCl, 1 mM DTT, and 10% glycerol). After a 10 min incubation at room temperature, fluorescence polarization was read using a Tecan SPARK fluorescence plate reader, and binding data were analyzed with Graphpad Prism version 10 using a single-site binding model.

### Yeast two-hybrid assays

For yeast two-hybrid assays with *A. thaliana* proteins, *A. thaliana* ASY3 fragments were PCR-amplified and cloned into the *NdeI* site of pGADT7 (Clontech # 630442) via isothermal assembly to generate activation domain (AD) fusions. The SMC1(head) construct (residues 1-181 and 1030-1218 of *A. thaliana* SMC1 fused by a GSGSASAG linker) was synthesized by IDT and cloned into the *EcoRI* site of pBridge (Clontech # 630404) via isothermal assembly to generate a binding domain (BD) fusion. Subsequently, the REC8 C-WHD domain (residues 551-627) was synthesized by IDT and cloned into the *BglII* site of the same vector.

For yeast two-hybrid assays with *M. musculus* proteins, *M. musculus* SYCP2 fragments were PCR-amplified and cloned into the *NdeI* site of pGADT7 (Clontech # 630442) via isothermal assembly to generate activation domain (AD) fusions. The SMC1(head) construct (residues 2-169 and 1056-1217 of *M. musculus* SMC1β fused by a GSGSASAG linker) was synthesized by IDT and cloned into the *EcoRI* site of pBridge (Clontech # 630404) via isothermal assembly to generate a binding domain (BD) fusion. Subsequently, the RAD21L C-WHD domain (residues 457-550) or REC8 C-WHD domain (residues 521-591) was synthesized by IDT and cloned into the *BglII* site of the same vector.

pBridge (BD) and pGADT7 (AD) plasmids were transformed into *S. cerevisiae* strains Y187 (*MATα*) and AH109 (*MATa*), respectively, and transformants were selected on CSM -TRP (for BD plasmids) and -LEU (for AD plasmids) agar plates at 30°C for 3-5 days. Haploid strains were mated on YPD agar, and diploids were selected on CSM -LEU -TRP (-LT) agar plates at 30°C for 3-5 days. Diploid cultures were grown overnight in 1.5 mL CSM -LT liquid media at 30°C in a tabletop Eppendorf Thermomixer until the OD_600_ reached ∼0.6-0.8. Cultures were adjusted to an OD_600_ of 0.8, and 4 μL of each culture was spotted on CSM -LT (no selection), CSM -LEU -TRP-HIS (-LTH; selection), and CSM -LEU -TRP -HIS -ADE (-LTHA; stringent selection) agar plates. Plates were incubated at 30°C for 3-5 days, and growth on -LTH and -LTHA plates was used as a readout for protein-protein interactions.

### ChIP-Seq

Chromatin immunoprecipitation sequencing (ChIP-seq) samples were prepared from 25 mL of meiotic cultures, 3 h after meiotic induction. Each sample was immediately cross-linked with 1% formaldehyde for 30 min at room temperature and then the formaldehyde was quenched by adding glycine to a final concentration of 125 mM. Fixed cells were collected by centrifugation at 2000 RPM for 3 min, the supernatant was removed, and cell pellets were immediately frozen at −80 °C until the ChIP-seq protocol was continued. Cell pellets were resuspended in 500 µL of lysis buffer (50 mM HEPES/KOH pH 7.5, 140 mM NaCl, 1 mM EDTA, 1% Triton X-100, 0.1% sodium deoxycholate) with protease inhibitors (1 mM PMSF, 1 mM Benzamidine, 1 mg/ml Bacitracin, one Roche Tablet (catalog # 11836170001) in 10 ml) and glass beads in a biopulveriser. Samples were sonicated at 15% amplitude for 5×15 s to obtain DNA at an average length of 500 bp. Sonicated cell lysate was centrifuged at 14,000 RPM for 10 min at 4 °C to remove cell debris, and the supernatant containing soluble chromatin fraction was isolated. In total, 50 µL of the chromatin fraction was set aside from each sample to use as input samples. To the remaining chromatin fraction, we added 2 µL of anti-Red1 antibody (Lot #16440; kind gift from Nancy Hollingsworth) and incubated rotating overnight at 4 °C. Antibodies and associated DNA were then isolated using Gammabind G Sepharose beads (GE Healthcare Bio, catalog # 17-0885-01). Crosslinks were reversed by incubating samples and input at 65 °C for at least 6 h. Proteins and RNAs were removed using proteinase K and RNaseA. Libraries for sequencing were prepared by PCR amplification using Illumina TruSeq DNA sample preparation kits v1 and v2. Libraries were quality checked on 2200 Tapestation. Libraries were quantified using Qubit analysis prior to pooling. The ChIP libraries were sequenced on Element Biosciences AVITI sequencer at the NYU Biology Genomics core to yield 150 bp paired-end reads. For spike-in normalization (SNP-ChIP), SK288c cross-linked meiotic samples were added to respective samples at 20% prior to ChIP processing (Vale-Silva *et al*, 2019).

SNP-ChIP libraries were sequenced on Biosciences AVITI sequencer to yield 150 bp paired-end reads.

### Processing of reads from Illumina sequencing for ChIP-seq analyses

Illumina output reads were processed in the following manner. The reads were mapped to the SK1 genome (GCA_002057885.1) using BWA (Yue *et al*, 2017). SNP-ChIP library reads were aligned to concatenated genome assemblies of SK1 and S288c genomes (Vale-Silva *et al*, 2019). Only reads that matched perfectly to the reference genome were retrieved for further analysis. 3ʹ ends of the reads were extended to a final length of 200 bp using MACS3 and probabilistically determined PCR duplicates were removed. The input and ChIP pileups were SPMR-normalized (single per million reads) and fold-enrichment of ChIP over input data was used for further analysis.

ChIP-seq datasets were made up of at least two biological replicates that were merged prior to processing. All datasets were normalized to the global mean of one and regional enrichment was calculated.

### Chromosome spreads

Meiotic cells were collected at hour 2, 3, and 4 after meiotic induction and treated with 200 mM Tris pH 7.5/20 mM DTT for 2 min at room temperature and then spheroplasted in 2% potassium acetate/1 M Sorbitol/ 0.13 μg/μL zymolyase T100 at 30 °C. The spheroplasts were rinsed and resuspended in ice-cold 0.1 M MES pH 6.4/1 mM EDTA/0.5 mM MgCl_2_/1 M Sorbitol. Cells were placed on clean glass slides (soaked in ethanol and air-dried) and one volume of fixative (1% paraformaldehyde/3.4% sucrose/0.14% Triton X-100) was added to the cells and distributed across the slide by gentle tilting. Four volumes of 1% Lipsol were added to the slide and mixed by tilting. Cells were spread by a clean glass rod. Four volumes of fixative solution were added and slides were left to dry overnight and stored at −80 °C until the next day. Prior to staining, slides were washed in 1× PBS/0.4% Kodak Photoflo, and blocked with 1× PBS/1% chicken egg white albumin (Sigma Aldrich #A5503) on a shaker for 15 min at room temperature. Slides were hybridized with primary antibodies at 37 °C for 1 h, washed with 1× PBS/0.05% Triton-X three times for 5 min each, and hybridized with secondary antibodies at 37 °C for 1 h. Slides were washed once with 1× PBS/0.05% Triton-X, once with 1× PBS/0.4% Kodak Photoflo, and twice with dH_2_0/0.4% Kodak Photoflo and air-dried prior to mounting with VECTASIELD with DAPI mounting medium (VWR #H-1200-10) and a VWR Micro Cover Glass, No. 1; 60 × 24 mm (VWR #48393 106). Red1 was detected using anti-Red1 rabbit serum (Lot #16441) at 1:200 and Alexa Fluor 488 anti-rabbit antibody at 1:2000. Gmc2 was detected using anti-Gmc2 mouse serum at 1:500 (gift from Amy MacQueen), and Alexa Fluor 594 anti-mouse at 1:2000. Stained slides were stored at 4 °C prior to imaging.

### Microscopy and cytological analysis

Images were collected on a Deltavision Elite imaging system (GE) equipped with an Olympus 100×/1.40 NA UPLSAPO PSF oil immersion lens and an InsightSSI Solid State Illumination module. Images were captured using an Evolve 512 EMCCD camera in the conventional mode and analyzed using ImageJ software. Box-and-whisker plots were generated in Microsoft Excel.

### Western blot analysis

For protein extraction, 5 mL of synchronous meiotic culture were harvested at T = 4 hours. Cells were immediately resuspended in 5 mL 5% trichloroacetic acid and incubated on ice for at least 10 min. Cells were collected by centrifugation, washed with 1 mL acetone, transferred to FastPrep tubes, and air dried overnight. 150 μL Tris-EDTA/250 mM DTT and glass beads were added to the cell pellet and cells ruptured in a bead beater. pH was adjusted by adding 20 μL of 1 M Tris base. SDS loading dye was added prior to boiling samples for 4 min. In all, 10 μL of protein sample were loaded onto an 8% polyacrylamide gel. Protein samples were resolved and transferred to a PVDF membrane using a Mini Trans-Blot Cell (BioRad). Membranes were blocked with 5% milk/PBST solution and then hybridized to rabbit anti-Red1 antibody (Lot #16440; kind gift from Nancy Hollingsworth) or rabbit anti-H2A antibody (Abcam, ab13923) as loading control. Anti-Rabbit-HRP from Kindle Biosciences (R1005) was used with 1-shot Digital-ECL substrate (R1003) to visualize protein. Images were taken with a digital camera.

## Data Availability

The ChIP-Seq data sets produced in this study have been deposited with the Gene Expression Omnibus under accession number GSE338394 (https://www.ncbi.nlm.nih.gov/geo/query/acc.cgi?acc=GSE338394).

The pipeline used to process SNP-ChIP reads and calculate spike-in normalization factor can be found at (https://github.com/hochwagenlab/ChIPseq_functions/tree/master/ChIPseq_Pipeline_hybrid_ge nome). R functions used for data processing can be found at (https://github.com/hochwagenlab/hwglabr2).

## Author Contributions

F.M.D. performed yeast genetics, Alphafold structure predictions, and biochemical reconstitution of protein-protein interactions, prepared figures, and wrote the manuscript with K.D.C.. C.R.M. performed synchronous meiosis, meiotic chromosome spreads, and ChIP-Seq data collection and analysis. J.H. performed yeast two-hybrid assays and biochemical reconstitution of protein-protein interactions. A.H. provided resources, guided experimental approaches, prepared figures, and contributed to writing the manuscript. K.D.C. provided resources, guided experimental approaches, prepared figures, and wrote the manuscript with F.M.D..

## Acknowledgements

The authors thank members of the Corbett and Hochwagen labs for helpful discussions. This work was supported by the National Institutes of Health (R35 GM144121 to KDC; R35 GM148223 to A.H.). F.M.D. was supported by the UCSD CCSC Training Grant (NIH T32 CA009523).

---

**Figure S1.**
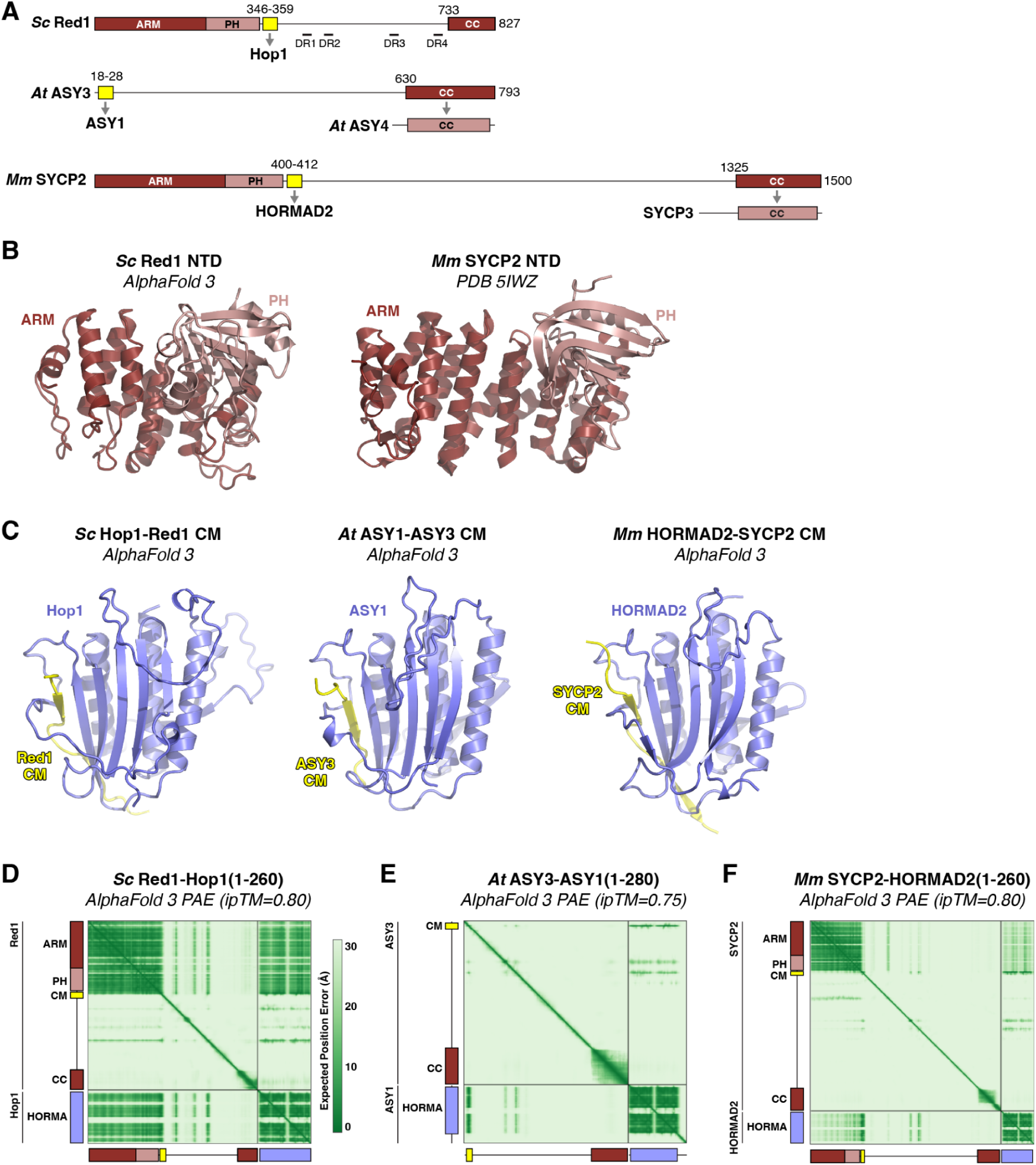
Architecture of meiotic chromosome axis core proteins. (A) Domain schematic of *S. cerevisiae* Red1 (Sc Red1), *A. thaliana* ASY3 (At ASY3), and *M. musculus* SYCP2 (Mm SYCP2). ARM: armadillo repeat domain; PH: pleckstrin homology domain; CM: closure motif; CC: coiled-coil. Gray arrows indicate known interactions between closure motifs and HORMAD binding partners, and between coiled-coil domains and axis core protein binding partners (West *et al*, 2019). (B) Axis core protein N-terminal domain structures from *S. cerevisiae* Red1 (left; AlphaFold 3 predicted) and *M. musculus* SYCP2 (right; PDB ID 5IWZ) (Feng *et al*, 2017). (C) AlphaFold 3 predicted structures of axis core protein closure motif-HORMA complexes from *S. cerevisiae* (left), *A. thaliana* (middle), and *M. musculus* (right). Closure motifs are shown in yellow, and HORMA domains are shown in blue. (D) AlphaFold 3 predicted aligned error (PAE) plot for *S. cerevisiae* Red1 and Hop1 (residues 1-260). (E) AlphaFold 3 predicted aligned error (PAE) plot for *A thaliana* ASY3 and ASY1 (residues 1-280). (F) AlphaFold 3 predicted aligned error (PAE) plot for *M. musculus* SYCP2 and HORMAD2 (residues 1-260). AlphaFold structure predictions with SYCP2 and HORMAD1 do not unambiguously identify a HORMAD1-binding closure motif in SYCP2 (not shown).

**Figure S2.**
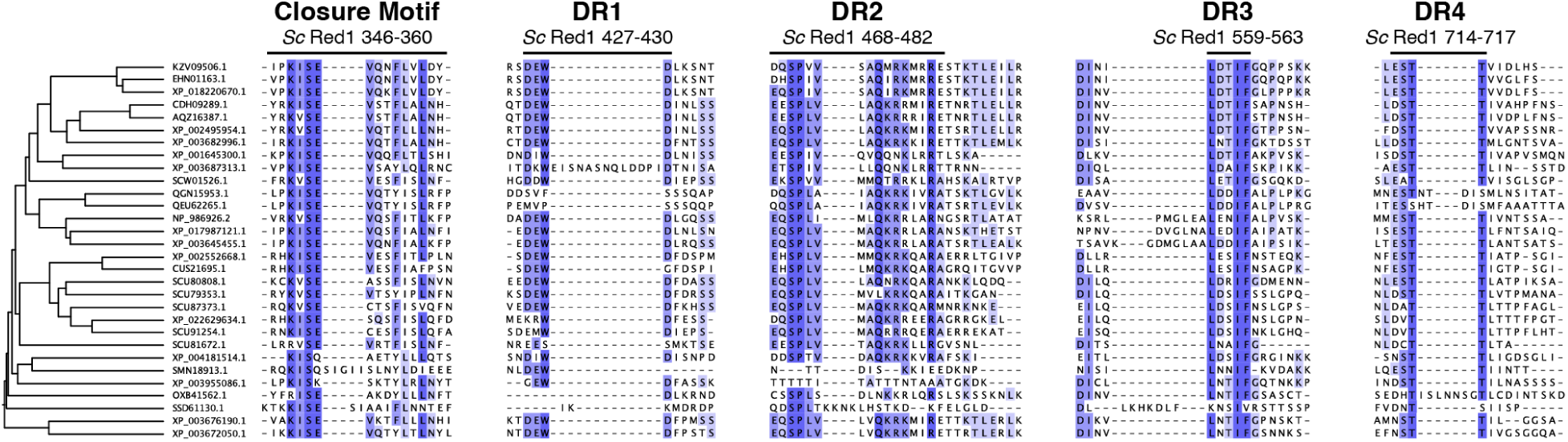
Conserved motifs in the budding yeast Red1 disordered region. Sequence alignment of 30 budding-yeast Red1 homologs, with NCBI accession numbers noted and average-distance tree calculated by Jalview (Troshin *et al*, 2011). The closure motif and DR1 through DR4 are noted.

**Figure S3.**
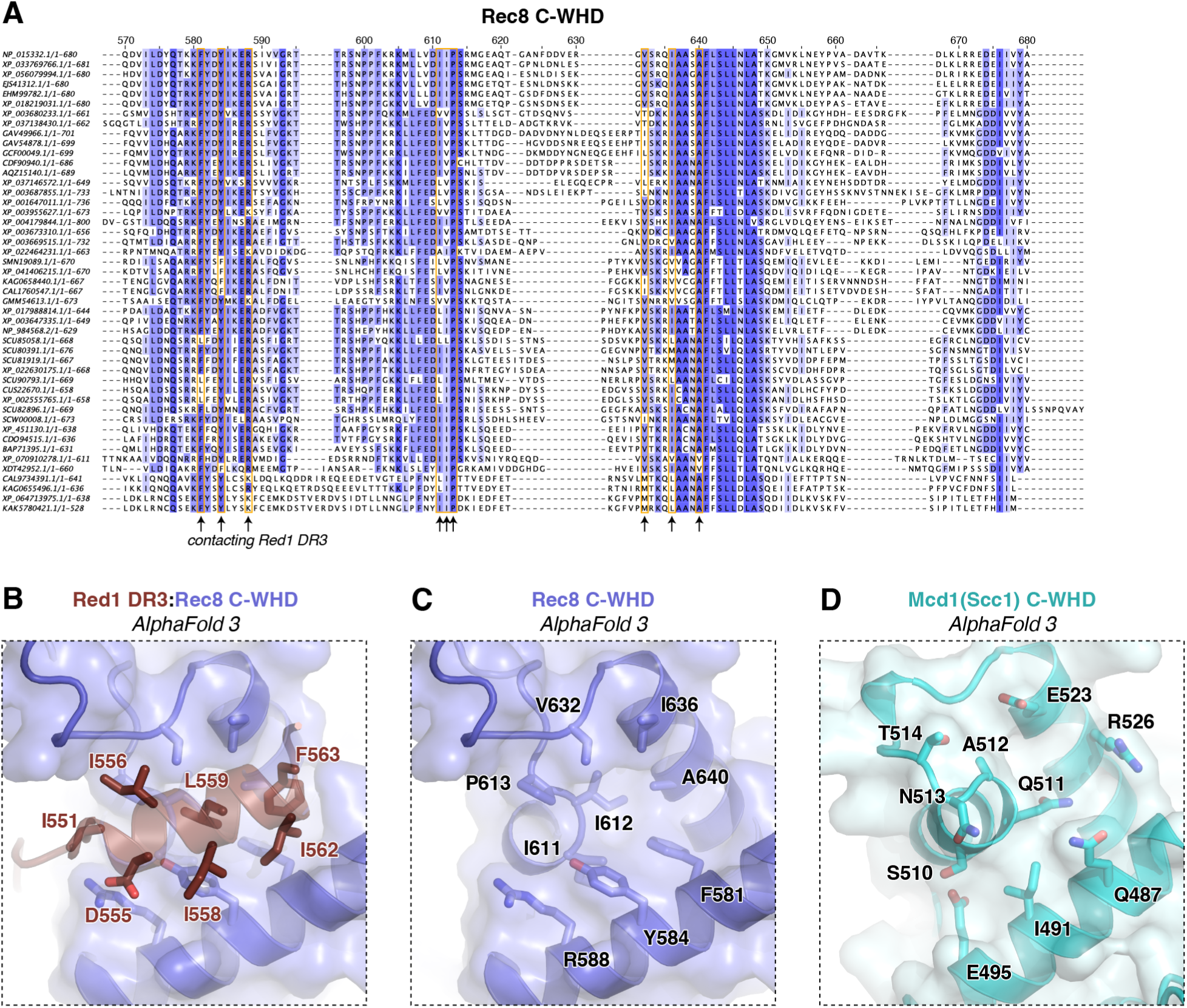
The Rec8 C-WHD Red1-binding surface is conserved in fungi. (A) Sequence alignment of fungal Rec8 C-WHD domains. Numbering shown is for *S. cerevisiae* Rec8 (top line). Residues predicted by AlphaFold to contact Red1 DR3 are boxed in orange and indicated with arrows at bottom. (B) AlphaFold 3 structure prediction of the Rec8 C-WHD domain (blue) bound to the Red1 DR3 motif (brown). Contacting residues on both partners are shown in sticks, and Red1 residues are labeled. (C) View of the Rec8 C-WHD domain as in panel (B) with Red1 DR3 not shown, and Rec8 residues labeled. (D) View of the Mcd1 (Scc1) C-WHD (AlphaFold 3 prediction) equivalent to the view in panel (C).

**Figure S4.**
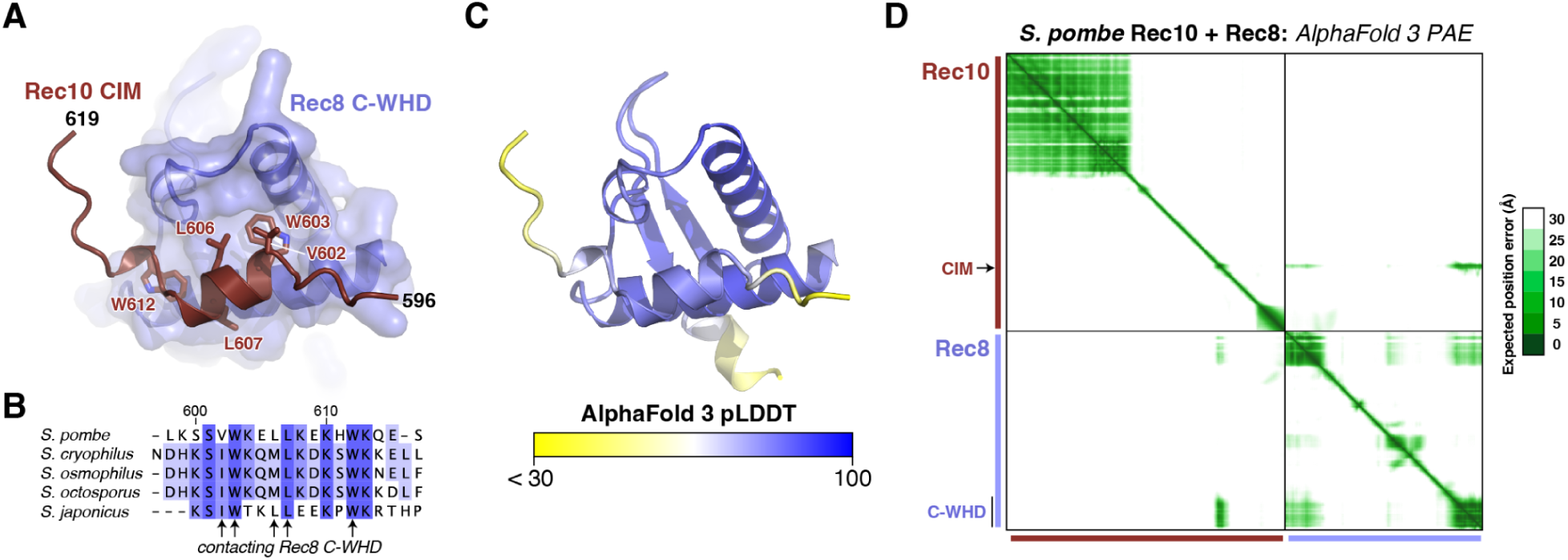
*S. pombe* Rec10 is predicted to bind the Rec8 C-WHD. (A) Cartoon view of an AlphaFold 3 predicted structure of *S. pombe* Rec10 (homolog of *S. cerevisiae* Red1; brown) binding to the Rec8 C-WHD (blue). Residues in the putative Rec10 CIM that are predicted to contact Rec8 are shown as sticks and labeled. (B) Sequence alignment of the putative CIM region of *S. pombe* Rec10. Sequences shown are NCBI NP_594524.1 (*Schizosaccharomyces pombe*), XP_013023355.1 (*Schizosaccharomyces cryophilus*), XP_056038338.1 (*Schizosaccharomyces osmophilus*), XP_013020446.1 (*Schizosaccharomyces octosporus*), and XP_002175829.1 (*Schizosaccharomyces japonicus*). (C) View equivalent to panel (A), colored by local confidence (pLDDT). (D) AlphaFold 3 predicted aligned error (PAE) plot for the predicted structure shown in panel (A) (full-length sequences for both proteins were used in the prediction). The overall AlphaFold 3 ipTM (interface predicted template modeling) score for this prediction is 0.27.

**Figure S5.**
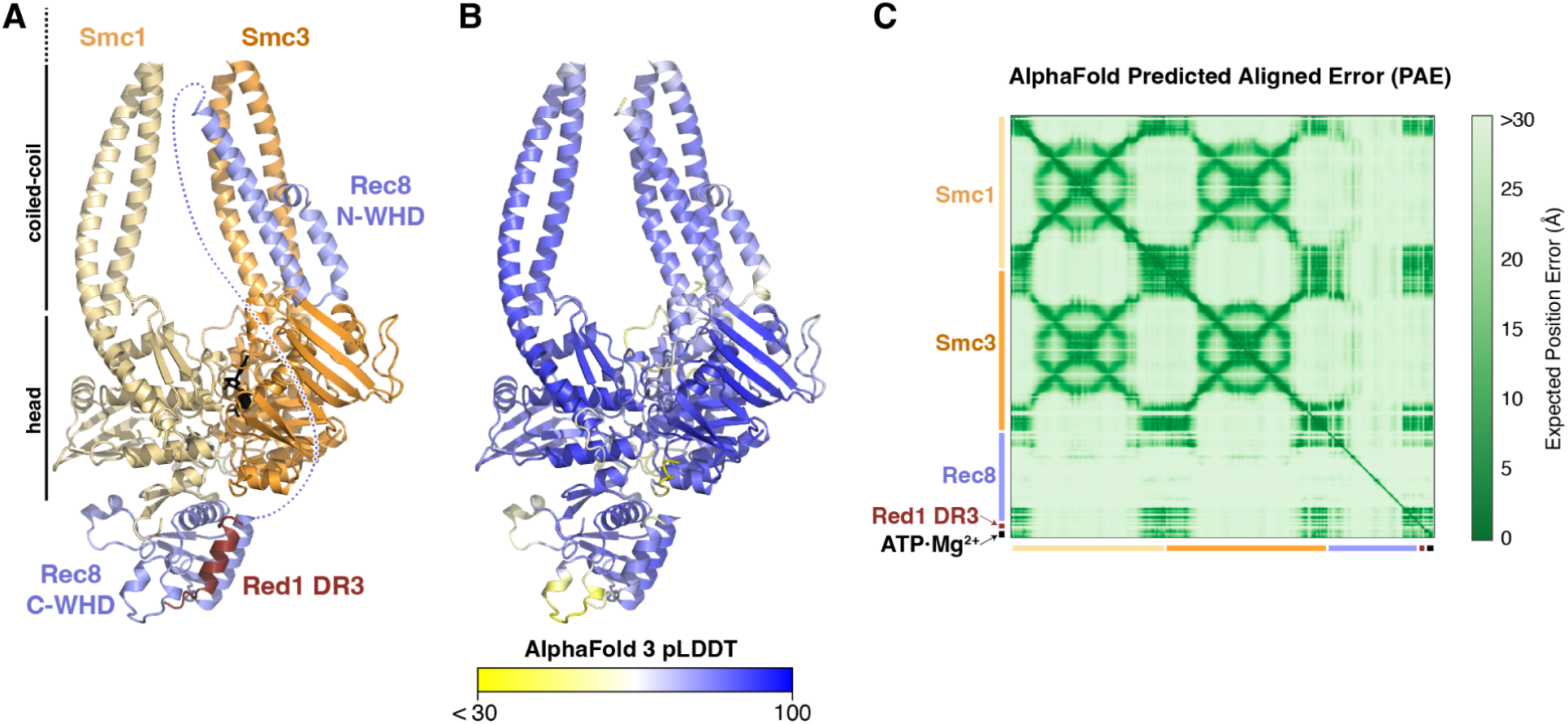
Predicted structure of the *S. cerevisiae* meiotic cohesin complex bound to Red1 DR3. (A) Cartoon view of an AlphaFold 3 predicted structure of *S. cerevisiae* Smc1 (light yellow), Smc3 (orange), Rec8 (light blue; central disordered region shown as dotted line), Red1 DR3 (brown), and ATP·Mg^2+^ (black). The extended coiled-coil and hinge domains of Smc1 and Smc3 are not shown. (B) View equivalent to panel (A), colored by local confidence (pLDDT). (C) AlphaFold 3 predicted aligned error (PAE) plot for the predicted structure shown in panel (A). The overall AlphaFold 3 ipTM (interface predicted template modeling) score for this prediction is 0.46.

**Figure S6.**
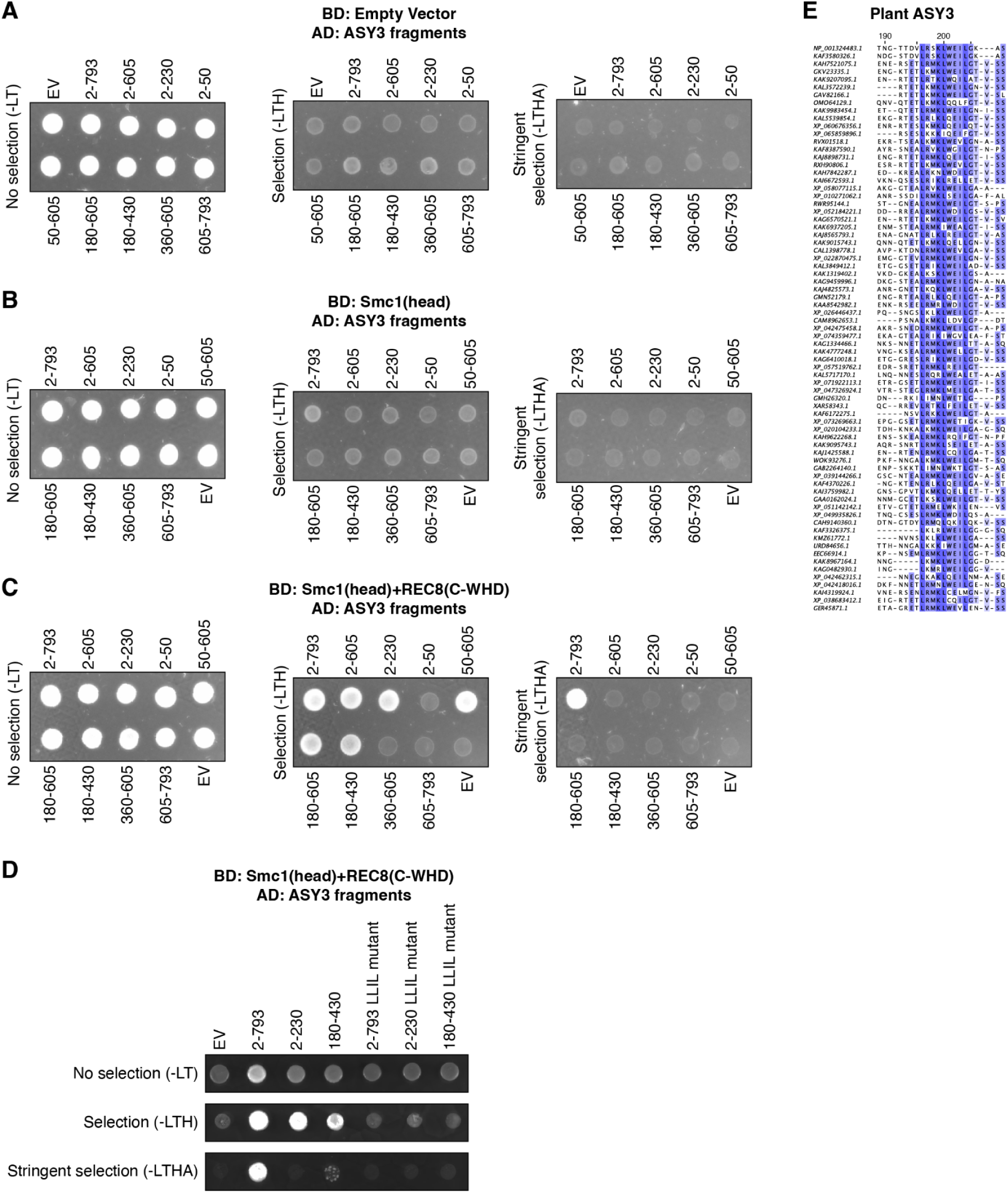
Identification of a cohesin binding motif in plant ASY3. (A) Yeast two-hybrid assay using empty binding-domain (BD) plasmid and AD plasmids with *A. thaliana* ASY3 fragments as indicated. EV: empty vector. No selection (-LT): non-selective media lacking leucine and tryptophan; Selection (-LTH): media further lacking histidine; Stringent selection (-LTHA): media further lacking adenine. (B) Yeast two hybrid assay using *A. thaliana* SMC1(head) BD plasmid and AD plasmids with *A. thaliana* ASY3 fragments as indicated. (C) Yeast two hybrid assay using *A. thaliana* SMC1(head) + REC8(C-WHD) BD plasmid and AD plasmids with *A. thaliana* ASY3 fragments as indicated. (D) Yeast two hybrid assay using *A. thaliana* SMC1(head) + REC8(C-WHD) BD plasmid and AD plasmids with *A. thaliana* ASY3 fragments as indicated. LLIL mutant: ASY3 L196D, L200D, I203N, L204E). (E) Section of a multiple sequence alignment of plant ASY3 proteins around the putative REC8(C-WHD) binding motif. Residue numbering is for *A. thaliana* ASY3 (top line).

**Figure S7.**
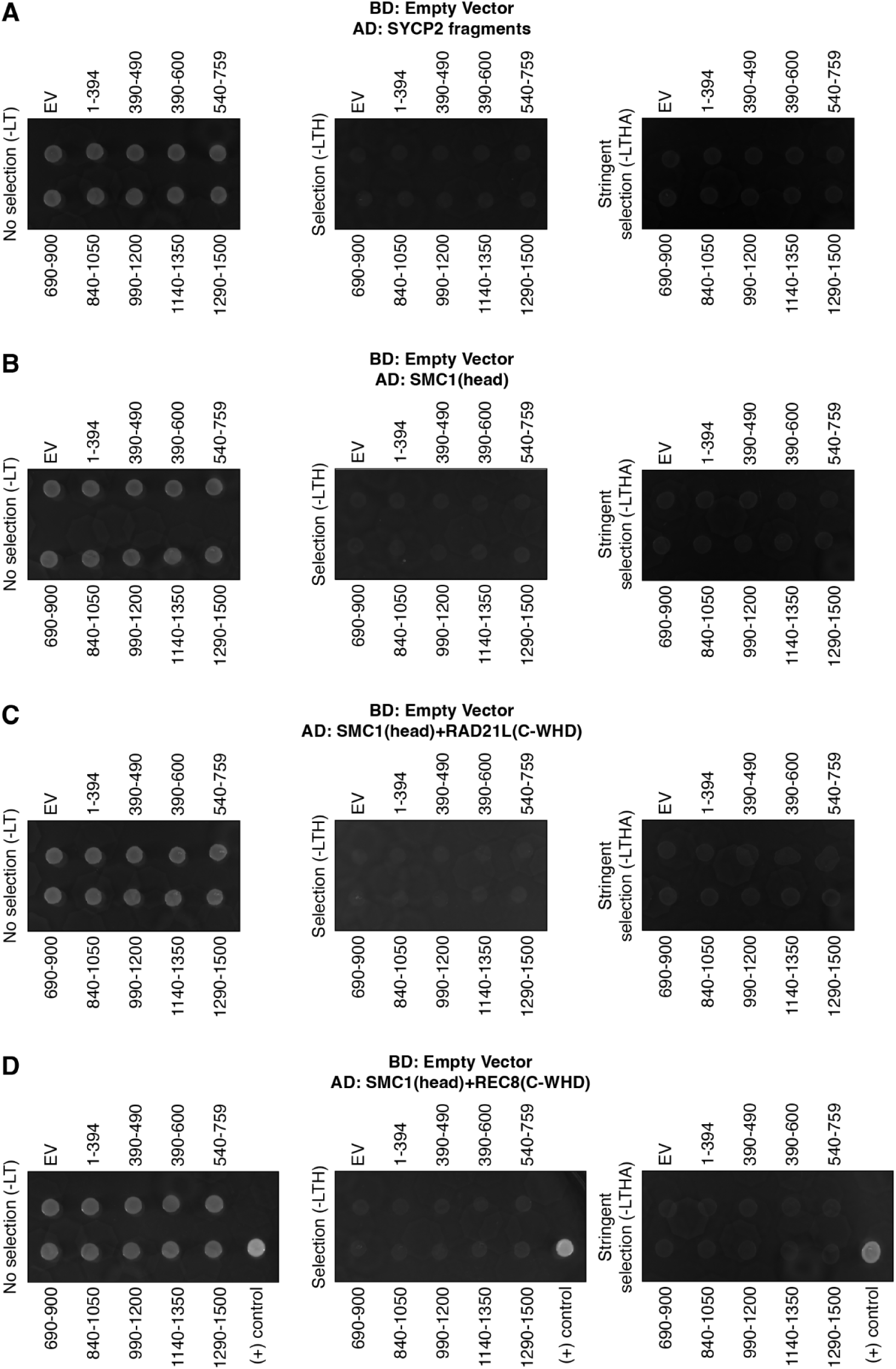
Yeast two hybrid assays with *M. musculus* axis proteins. (A) Yeast two-hybrid assay using empty binding-domain (BD) plasmid and AD plasmids with *M. musculus* SYCP2 fragments as indicated. EV: empty vector. No selection (-LT): non-selective media lacking leucine and tryptophan; Selection (-LTH): media further lacking histidine; Stringent selection (-LTHA): media further lacking adenine. (B) Yeast two hybrid assay using *M. musculus* SMC1(head) BD plasmid and AD plasmids with *M. musculus* SYCP2 fragments as indicated. (C) Yeast two hybrid assay using *M. musculus* SMC1(head)+RAD21L(C-WHD) BD plasmid and AD plasmids with *M. musculus* SYCP2 fragments as indicated. (D) Yeast two hybrid assay using *M. musculus* SMC1(head)+REC8(C-WHD) BD plasmid and AD plasmids with *M. musculus* SYCP2 fragments as indicated. “(+) control” indicates a BD-p53+AD-SV40 large T antigen positive control.

